# The modified RNA base acp3U is an attachment site for N-glycans in glycoRNA

**DOI:** 10.1101/2023.11.06.565735

**Authors:** Yixuan Xie, Helena Hemberger, Nicholas A. Till, Peiyuan Chai, Christopher P. Watkins, Charlotta G. Lebedenko, Reese M. Caldwell, Benson M. George, Carolyn R. Bertozzi, Benjamin A. Garcia, Ryan A. Flynn

## Abstract

We recently identified glycoRNA—a previously undescribed glycoconjugate—which consists of RNAs modified with secretory N-glycans and presented on the cell surface. While previous work supported a covalent linkage between RNA and glycans, the direct chemical nature of the RNA-glycan connection was not described. Here we develop a sensitive and scalable protocol to detect and characterize native glycoRNAs. Leveraging periodate oxidation and aldehyde ligation (rPAL) and Sequential Window Acquisition of all Theoretical Mass Spectra (SWATH-MS), we identified the modified RNA base 3-(3-amino-3-carboxypropyl)uridine (acp3U) as a site of attachment of N-glycans in glycoRNA. The sensitivity and robustness of rPAL provided the first evidence of a direct glycan-RNA linkage, and its flexibility will enable further characterization of glycoRNA biology.

## Introduction

We recently showed that a sialic acid-specific metabolic chemical reporter (MCR), N-azidoacetylmannosamine- tetraacylated (Ac_4_ManNAz), is conjugated to small noncoding RNAs in mammalian cells through N-glycan linkages(*1*). These sialylated glycoRNAs (sialoglycoRNAs) can be isolated from a variety of cell types, grown in culture or live animals, and critically, can be presented on the outer surface of the cell membrane(*1*). Missing from this work was a detailed characterization of the chemical linkage that enables RNA glycosylation. To address this missing link (**Fig. 1A**), we developed a highly sensitive and scalable chemical strategy to label sialoglycoRNAs and provide evidence that acp3U is a direct linker between mammalian small RNAs and N-glycans.

**Fig. 1.**
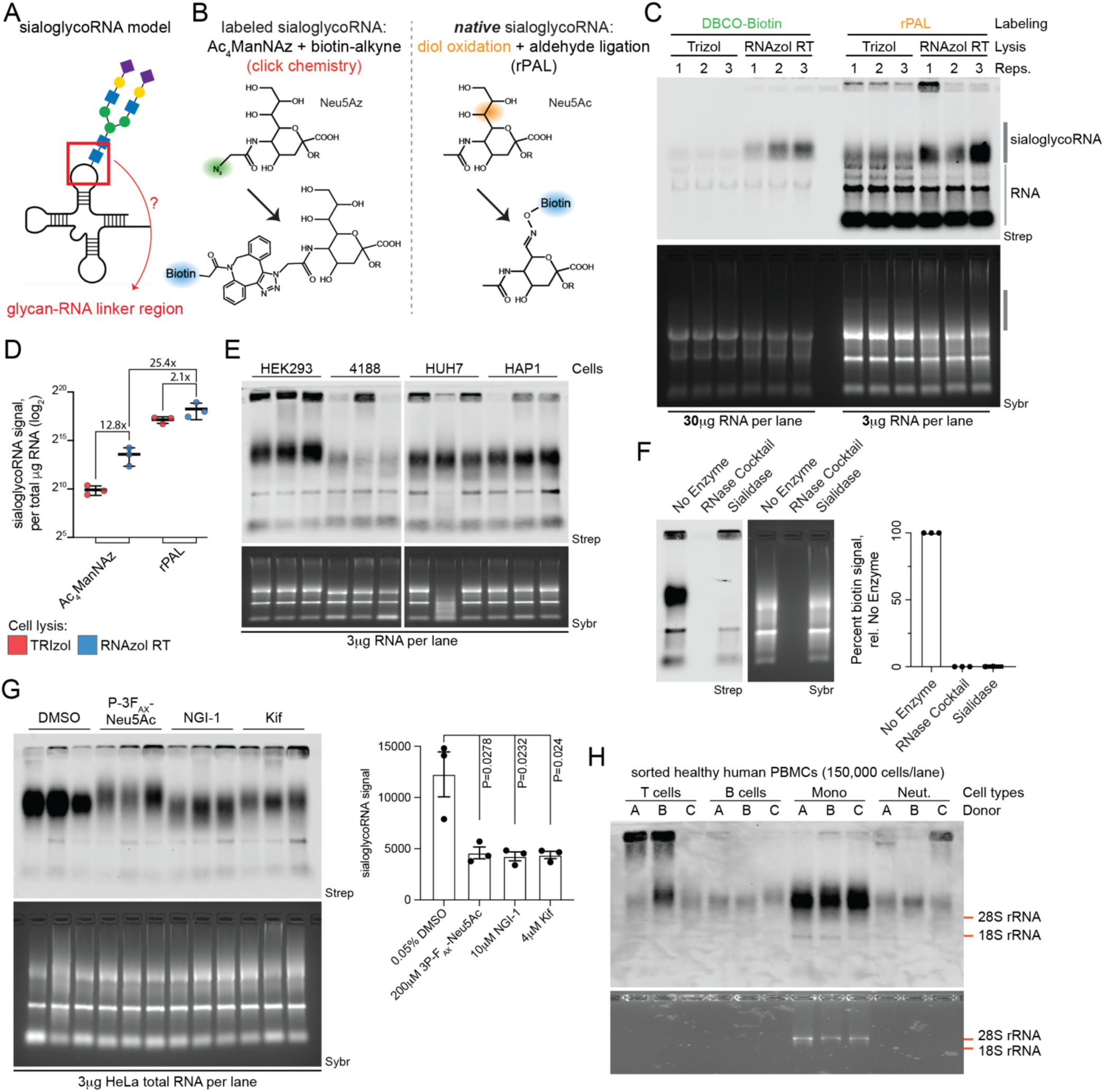
Selective oxidation and aldehyde labeling enables sensitive detection of native sialoglycoRNAs. (**A**) Cartoon of a sialoglycoRNA highlighting the glycan-RNA linker region which has not been previously defined. (**B**) Schematic of sialic acid labeled strategies. Left: after uptake and conversion of Ac_4_ManNAz into Neu5Az, copper-free click chemistry can selectively label the Neu5Az with a biotin handle for detection. Right: native sialic acid diols can be oxidized and the resulting aldehyde ligated to a biotin reagent for detection (rPAL). (**C**) RNA blotting of HeLa total RNA labeled and detected with Ac_4_ManNAz paired with copper-free click of Dibenzocyclooctyne-PEG4-biotin (DBCO-biotin) or native sialic acids paired with rPAL labeling. Total RNA was extracted with either TRIzol or RNAzol RT to compare relative efficiencies of sialoglycoRNA detection. In gel detection of total RNA with SybrGold (Sybr, bottom) and on membrane detection of biotin (Streptavidin-IR800, top) is shown. The region labeled “sialoglycoRNA” was quantified. (**D**) Quantification of data in (C). Each datapoint (biological triplicate) is displayed with the standard error of the mean (SEM). (**E**) RNA blotting of HEK293, 4188, HUH7, and HAP1 total RNA. Detection of the resulting sialoglycoRNA signal was accomplished by rPAL labeling and quantified to the right. Sybr and Strep detection is as in C. (**F**) RNA blotting of HeLa total RNA that was treated in vitro with no enzyme, an RNase A and RNase T1 cocktail, or Sialidase. Detection of the resulting sialoglycoRNA signal was accomplished by rPAL labeling and quantified to the right. Sybr and Strep detection is as in C. Each datapoint (biological triplicate) is displayed with the SEM. (**G**) RNA blotting of total RNA from HeLa cells treated with various chemical inhibitors. Inhibitors were added to cells in complete media for 24 hours: 0.05% DMSO, 200 μM P-3_FAX_-Neu5Ac, 10 μM NGI-1, or 4 μM Kifensuesine, after which RNA was collected and processed for rPAL labeling. Sybr and Strep detection is as in A. SialoglycoRNA signal was accomplished by rPAL labeling and quantified to the right; each datapoint (biological triplicate) is displayed with the SEM and statistical analysis was performed using an unpaired t-test. (**H**) RNA blotting of total RNA from four sorted populations of human PBMCs including CD19 (B cells), CD3 (T cells), CD14 (monocytes and macrophages), and CD16 (NK cells, monocytes, and neutrophils) positive cell RNA from approximately 150,000 cells were used for each lane across the four populations.

### Native sialoglycoRNA detection

Our previous work relied on metabolic conversion of an azide-modified sialic acid precursor, peracetylated *N*-azidoacetylmannosamine (Ac_4_ManNAz) to the corresponding azido sialic acid and its biosynthetic incorporation into the N-glycans of glycoRNA (**Fig. 1B**). However, the reliance of this method on cellular uptake and metabolism made large scale preparations of glycoRNA difficult, thus limiting our ability to perform detailed structural analyses by mass spectrometry. As well, incorporation of unnatural azidosugars into glycans may have perturbing effects on their natural trafficking mechanisms and bioactivity. We therefore pursued an alternative approach to label and enrich glycoRNAs that leverages the periodate-mediated oxidation of vicinal diols to aldehydes and their subsequent ligation to amine-containing reagents or solid supports. This method has been used widely to characterize sialoglycolipids and proteins(*2–6*). Paired with aminooxy containing molecules, this reaction forms a stable oxime bond without requiring further derivatization. While this approach is reasonably selective for sialic acid-containing glycans, the presence of a 2’, 3’ vicinal diol at the 3’ terminal ribose of RNA poses a challenge to its application to glycoRNA. While the 7’, 8’ diol in sialic acid is quite reactive—and thus can be derivatized at physiological pH with short reaction times(*6*)— complete oxidation of ribose diols in RNA requires more acidic buffers and longer reaction times(*7*). With this insight into the differential reactivity of sialic acid diols and ribose diols, we reasoned that in a mixture of RNA nucleotides and sialic acid sugars, mild oxidation conditions could achieve selective sialic acid diol labeling (**Fig. 1B**). We examined cell lysis methods,RNA extraction reagents (TRIzol vs. RNAzol RT), and specific precipitation parameters for RNA column clean-ups (**Supplemental Information**) to develop a robust protocol for high recovery of small RNA (**Fig. S1A-C**).

To maintain a simple procedure, we set out to develop a reaction scheme that would both oxidize diols and ligate newly generated aldehydes (periodate oxidation and aldehyde labeling, PAL) to a labeling reagent in the same reaction without purification. Conditions including pH, salt concentration, salt type, and temperature were screened using RNAzol RT-extracted HeLa cell total RNA (**Fig. S1D-F**). An RNA optimized PAL protocol (rPAL, **Supplemental Information**) includes a pre-blocking step with a free aldehyde reagent, which reduces background signal, as well as mucinase digestion (**Fig. S1G, S1H**) over 2.5 hours. Further development of the RNA Northern blot transfer buffer conditions (pH, salt, and time, **Fig. S2A-D**) resulted in enhanced transfer. Extracting RNA from Ac_4_ManNAz labeled HeLa cells with TRIzol and either performing copper-free click(*1*) or rPAL, we found that rPAL generates approximately 150x the amount of signal (**Fig. 1C, 1D**). We repeated this comparison from Ac_4_ManNAz-labeled cells and while there was not a significant difference in total RNA extracted with TRIzol vs RNAzol RT (**Fig. S1A**), we saw a 12.8x and 2.1x gain in glycoRNA signal with Ac_4_ManNAz and rPAL respectively (**Fig. 1D**). Integrating both the rPAL and RNAzol RT RNA extraction, we can achieve at least 25-fold increased signal recovery per mass of RNA compared to an updated Ac_4_ManNAz-strategy that uses RNAzol RT extractions (**Fig. 1D**). While rPAL improves sensitivity of apparent high molecular weight (MW) glycoRNA species, it also induces background labeling; most notably within the 18S rRNA and the small RNA pool (**Fig. 1D** and elsewhere). This is expected given that the 3’ ends of RNAs should (mostly) contain a 2’-3’ vicinal diol, which can also undergo periodate-based oxidation(*7*).

To demonstrate the flexibility of rPAL, we labeled total RNA stocks which were collected over 3 years ago from cells treated with Ac_4_ManNAz for 24 hours prior to collection. rPAL readily detects sialoglycoRNAs from the four archived RNA stocks sourced from HEK293, 4188, HUH7, and HAP1 cells (**Fig. 1E**). We can more directly interpret the levels of rPAL signal across each cell type due to differences in sialylation, while Ac_4_ManNAz-labeling requires consideration of metabolism and incorporation. Finally, to establish that the high MW signal is indeed sialoglycoRNA, we evaluated its sensitivity to enzymatic digestion. Incubation of purified RNA with RNase or sialidase and subsequent rPAL-labeling results in total loss of all rPAL signal after RNase but selective and complete loss of the high MW signal after sialidase treatment (**Fig. 1F**), consistent with rPAL-mediated sialic acid labeling of sialoglycoRNAs.

### rPAL characterization

To verify that the high MW signal is indeed small glycoRNAs, we used column-based separation of large and small RNAs; we found that the high MW rPAL signal, as well as the signal overlapping with the small RNA pool, is enriched with small RNAs but not the long RNAs (**Fig. S3A**). Cell fractionation demonstrated that the high MW signal co-purifies with crude cellular membrane extracts while it is depleted in the cytosolic fraction (**Fig. S3B, S3C**). Additionally, and in line with our reported topology of Ac_4_ManNAz-labeled sialoglycoRNAs, the rPAL-labeled high MW signal is sensitive to live-cell sialidase treatment. By treating live HeLa cells with vibrio cholerae (VC) sialidase, extracting RNA, and performing rPAL labeling, we could detect robust loss of sialoglycoRNA signal in as little as 15 minutes after addition of the VC-sialidase (**Fig. S3D**).

Because inhibition of N-glycosyltransferases resulted in loss of signal from Ac_4_ManNAz-labeled sialoglycoRNAs(*1*), we assessed how the rPAL signal from HeLa cells responded to treatment with the following inhibitors: 1) P-3F_AX_-Neu5Ac (sialic acid biosynthesis)(*8*); 2) NGI-1 (oligosaccharyltransferase)(*9*); and 3) Kifunensine (ɑ-mannosidase-I)(*10*). In all three cases, we observed loss of the rPAL-signal (**Fig. 1G**), consistent with our previous results detecting sialoglycoRNAs with Ac_4_ManNAz. P-3F_AX_-Neu5Ac and Kifunensine treatment, respectively, caused a larger shift in the apparent MW of the sialoglycoRNAs (**Fig. 1G**), which is in line with the effects seen with Ac_4_ManNAz. NGI-1 resulted in an apparent lower MW smearing of the signal (**Fig. 1G**), which was not previously seen with Ac_4_ManNAz.

To demonstrate one utility of the significantly increased sensitivity of rPAL and the ability to access samples where MCR-labeling would be challenging, we examined the sialoglycoRNAs from human peripheral blood mononuclear cells (PBMCs). We stained healthy human PBMCs with antibodies targeting CD19 (B cells), CD3 (T cells), CD14 (monocytes and macrophages), and CD16 (NK cells, monocytes, and neutrophils) and performed cell sorting (**Fig. S3E**). We sorted 150,000 cells per tube per donor (three total donors) after which RNA was extracted (total RNA amounts in the 100’s of nanograms per reaction) and processed with the rPAL labeling method. Across all four cell types sorted we can detect sialoglycoRNAs (**Fig. 1H**). All donors generated sialoglycoRNA signal from the four cell types analyzed, with the sorted CD14+ cells yielding the most total RNA as well as the strongest sialoglycoRNA signal (**Fig. 1H**). Some samples from each of the cell types also displayed ultrahigh MW signal (**Fig. 1H**), which is evident but less common in material isolated from cultured lines with higher cell numbers (e.g., **Fig. 2C**, lanes 1, 3, and 4). We note that minor MW change could be seen between donors of T cells (donor 3) and B cells (donor 2) (**Fig. 1H**). Taken together, these data demonstrate that rPAL labeling is highly sensitive and useful in low input material contexts. Given the robustness of the protocol, we predicted it would be easily scaled up for large enrichments useful for de novo discovery of the putative glycan-RNA linkage.

**Fig. 2.**
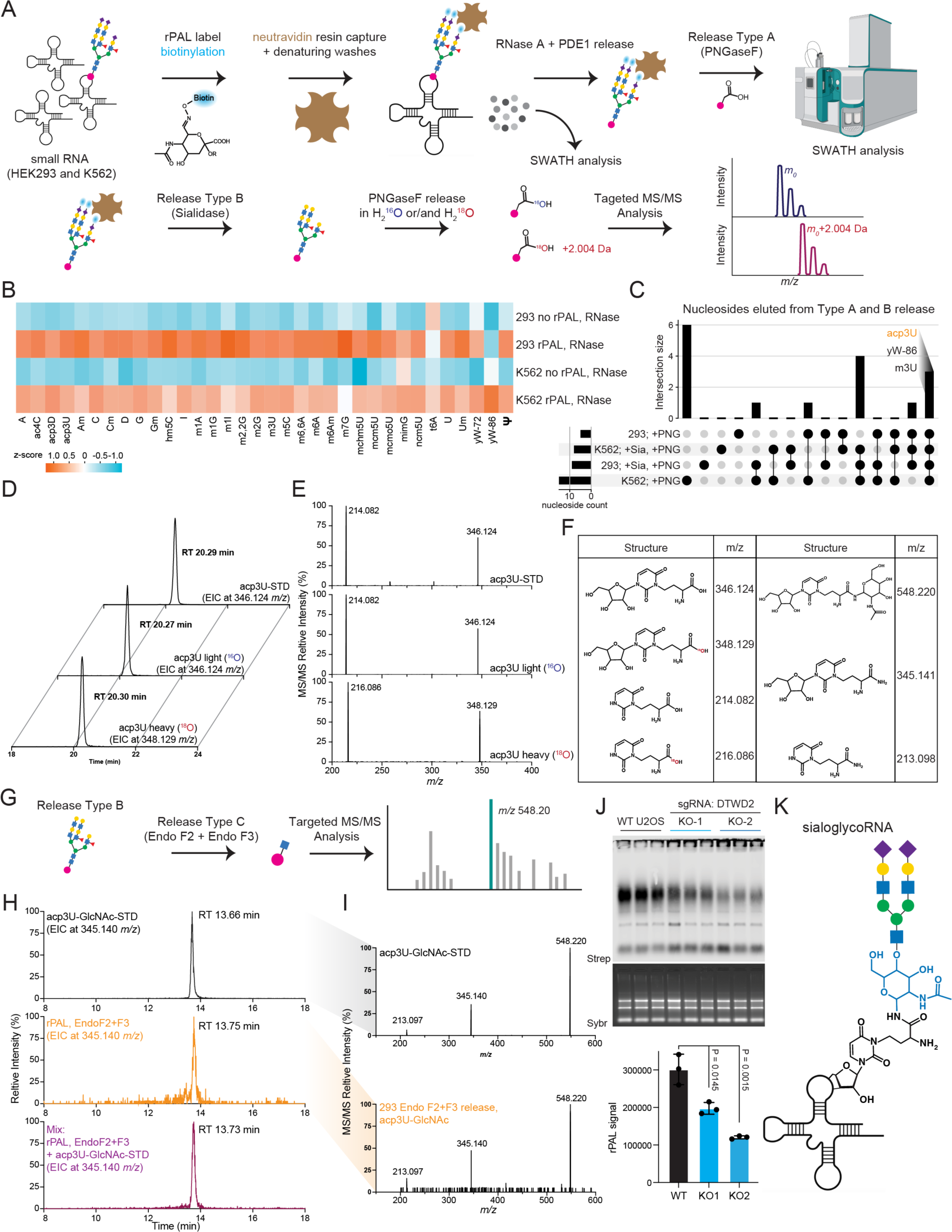
3-(3-Amino-3-carboxypropyl)uridine (acp3U) is an endogenous template for RNA glycosylation. (**A**) Schematic of the rPAL labeling, enrichment, RNase digestion, on-bead PNGaseF release, and SWATH-MS analysis (top, Release Type A). Orthogonal processing of post-RNase digested material with on-bead sialidase release followed by in solution PNGaseF digestion in the presence of heavy (^18^O) or light (^16^O) water (bottom). (**B**) Heatmap analysis of the nucleosides identified by SWATH-MS from HEK293 and K562 cells after the RNase digestion. Z-scores were calculated for each nucleoside between samples and colored from −1.0 to 1.0. (**C**) Upset plot intersection of the nucleosides identified by SWATH-MS from HEK293 and K562 cells from both the RNase digestion as well as the on-bead PNGaseF release experiments. Three nucleosides were found as present in all four datasets and are highlighted: acp3U, yW-86, m3U. (**D**) Extracted Ion Chromatogram (EIC, specific m/z shown) from the liquid chromatogram (LC) of acp3U-STD, Type B digestion with light (^16^O) water, and Type B digestion with heavy (^18^O) water. Retention times for each peak are annotated. (**E**) Tandem mass spectrum (MS/MS) analysis of the three peaks shown in (D). m/z values for the two major peaks in each trace are annotated. (**F**) Chemical structures of heavy and light acp3U, the ribose-loss fragmentation products, an acp3U-GlcNAc, an amidated form of acp3U (loss of GlcNAc), and ribose-loss of the amidated acp3U. (**G**) Schematic of the Release Type C: after on-bead sialidase (Release Type B) samples were further processed with Endoglycosidase F2 (Endo F2) and Endo F3, which would result in a GlcNAc scar (blue square) on the N-glycosylated molecule. (**H**) EIC (specific m/z shown) from the LC of acp3U-GlcNAc-STD, Endo F2+F3 released material from rPAL enriched RNA, and a mixture of these two samples (mix). Retention times for each peak are annotated. (**I**) MS/MS analysis of two peaks shown in (H). m/z values for the three major peaks in each trace are annotated. (**J**) rPAL blotting (top) and quantification (bottom) of total RNA samples extracted from U2OS wild type (WT) or two individual knockout clones of the DTWD2 gene. Each datapoint (biological triplicate) is displayed with the standard deviation. A t-test was performed to calculate the P values. (**K**) Final model of a sialoglycoRNA. The primary modification of RNA with acp3U provides a site upon which N-glycosylation can occur, directly connecting the GlcNAc of the chitobiose core to the rest of the N-glycan.

### Glyconucleoside identification with rPAL enrichment

In the broader context of RNA modifications, methylation and pseudouridylation are the most abundant RNA modifications; in particular, Watson-Crick face methylations (N-1-methyladenosine (m1A), N-3-methylcytosine (m3C), 5-methylcytosine (m5C), which are known for their dynamism(*11*). More chemically complex, but less abundant modifications are also well-represented in small RNAs, especially tRNAs(*12*). For tRNA and rRNA PTMs, we have nucleotide-scale resolution of a majority of RNA modifications(*11*, *12*). N-glycosylation represents a new epitranscriptomic modification; however our initial report lacked a direct characterization of the chemical linkage between a particular atom of RNA (base, ribose, or phosphate) and the chitobiose core of an N-glycan.

To address this, we coupled large scale rPAL labeling, high efficiency capture, biochemical purification, enzymatic release (Type A, B, and C below), and SWATH-MS(*13*) (**Fig. 2A**). Specifically we performed the rPAL labeling as described above and captured sialoglycoRNAs on neutravidin resin. After high temperature (45°C) and stringent (4M NaCl and 2M GuHCl) washes, RNA was hydrolyzed with a cocktail of nucleases. Released nucleotides from this step could be putatively from regions of rPAL-labeled RNAs that are surrounding the glyconucleoside. In total, we found 34 unique nucleosides eluted across the three elution types, including unmodified A, C, U, and G, from HEK293 and K562 cells (**Fig. 2B, Table S1**). Subsequently, to release nucleotides that are connected to N-glycan core structures we treated the resin with PNGaseF (**Fig. 2A**, Type A). To release intact glyconucleosides, we instead subjected the final material to a sialidase elution followed by off-bead PNGaseF treatment (**Fig. 2A**, Type B). We compared the abundance of each nucleoside between control and rPAL labeled RNA: we found that rPAL enriched RNA from HEK293 cells released 4 modified nucleosides from Type A release and 7 modified nucleosides from Type B releases, while rPAL enriched RNA from K562 cells released 15 and 7 modified nucleosides from Type A and B releases, respectively (**Fig. 2C, Table S1**). When examining the structures of these released nucleosides, we noticed a number of them contained carboxylic acid groups (7-aminocarboxypropylwyosine, yW-72; 7-aminocarboxypropyl-demethylwyosine, yW-86; 3-(3-amino-3-carboxypropyl)uridine, acp3U). This was of interest because the products of PNGaseF cleavage of N-glycoproteins are asparagine to aspartic acid diagnostic ‘scars’(*14*).

Of acp3U, yW-72, and yW-86, acp3U was most enriched across both HEK293 and K562 cells and in both Type A and B releases. Further, its synthesis has been described(*15*), can be incorporated into oligonucleotide synthesis(*16*), is found in the core region of bacterial and eukaryotic tRNAs(*17–19*), and has several functional roles(*20*, *21*). To confirm that our identification of the acp3U nucleoside was a result of PNGaseF cleavage and not contamination of an endogenous modified nucleoside, we repeated the PNGaseF digestion in the presence of heavy (H_2_^18^O) or light (H_2_^16^O) water from the sialidase eluted material, as previously developed for determining the glycosite on glycoproteins(*22*). We expect this would result in PNGaseF-released acp3U having a mass-shift of 2.004 *m/z* pair in MS1 level, while displaying another paired pyrimidine ion during the tandem MS/MS (**Fig. 2A, bottom**). Examination of the LC retention time of the heavy and light cleaved material demonstrated overlapping peaks at 20.3 min (**Fig. 2D**). MS/MS of these peaks revealed the expected parent masses and fragmentation patterns of heavy and light acp3U (**Fig. 2E**). To validate the identity of acp3U, we used a synthetic standard. Commercially available acp3U exists as a diastereomeric mixture consisting of the natural amino acid stereochemistry and the unnatural epimer (**Fig. S4**). Stereoisomerically pure acp3U was prepared from the corresponding enantiopure homoserine derivative through a Mitsunobu coupling step (**Supplemental Information**). This synthetic acp3U (acp3U-STD) exhibited an HPLC retention time matching that of the cell-derived material (**Fig. 2D, S5-13**). MS/MS fragmentation of the acp3U-STD, light, and heavy PNGaseF released material resulted in peaks as expected (**Fig. 4E, 4F**), with the heavy released material shifted by 2 Da. These data suggest that we do indeed observe PNGaseF-released acp3U from rPAL enriched RNA.

To obtain direct evidence of a glyconucleoside, we repeated the process as outlined in Fig. 2A (Type B release), however we performed an in vitro digestion with a cocktail of Endo F2 and Endo F3 (Type C release, **Fig. 2G**) to release N-glycoconjugates with first GlcNAc still connected to the modified polymer. We predicted that a carboxamide version of acp3U could serve as a template for N-glycosylation and thus synthesized an acp3U-GlcNAc standard (acp3U-GlcNAc-STD, **Fig. S14-19**) with a predicted mass of 548.20 *m/z* (**Fig. 2F, 2G**) to examine this hypothesis. We compared the elution profile of the acp3U-GlcNAc-STD to that of released material rPAL enriched RNA, eluted with EndoF2 and EndoF3 (**Fig. 2H**) and found co-eluting peaks at 13.66 and 13.75 minutes, respectively. MS/MS analysis of these peaks demonstrated the expected fragmentation patterns of acp3U-GlcNAc (**Fig. 2I, 2F**). To additionally confirm the elution profile of acp3U-GlcNAc from the RNA sample, we spiked in the acp3U-GlcNAc-STD material into the rPAL enriched Endo F2+F3 cleaved material. This “mix” sample showed the same elution profile as the rPAL enriched Endo F2+F3 cleaved material alone (**Fig. 2H**). To further investigate acp3U biosynthesis as the source for sialoglycoRNA production, we generated two knockout clones of the gene DTWD2 (**Fig. S20**), one of two enzymes that are responsible for generating cellular acp3U(*20*). We selected DTWD2 because it has been reported to exist both in the nucleus and cytosol(*20*) and thus could have activity on the most number of transcripts in a cell. Both DTWD2-KO clones resulted in a loss of rPAL signal as compared to WT U2OS cells (**Fig. 2J**), consistent with the idea that acp3U could serve as a chemical linker between an RNA transcript and an N-glycan.

## Discussion

Here we describe the identification of acp3U as an attachment site for N-glycans on tRNA (**Fig. 2K**), which was facilitated by the development of the periodate oxidation and aldehyde ligation (rPAL) method. rPAL coupled to SWATH-MS, enabled the identification of the first molecular linker between a N-glycan and an RNA, furthering the initial discovery of glycoRNAs(*1*) and laying a path for examining other chemical mechanisms of RNA glycosylation.

A series of carboxylic acid-containing nucleosides were eluted from rPAL-enriched RNA with PNGaseF including acp3U, yW-72, and yW-86; however, acp3U was most consistently observed and was quantitatively most enriched in the cell lines examined. Coupled to our genetic knockout data of the DTWD2 enzyme, our data strongly support a model where acp3U is one of the primary RNA modifications that enables cells to N-glycosylate RNA. We therefore propose a biosynthetic process for generation of cell surface sialoglyco-tRNAs bearing N-glycans on acp3U residues as shown in **Fig. 3**. In this model, tRNAs are biosynthesized in the nucleus and modified to create acp3U residues in the nucleus and/or cytosol, which is then converted to the carboxamide functionality by as-yet unknown enzymes. Translocation into the ER lumen would then situate the modified tRNA for modification by oligosaccharyltransferase (OST). Notably, Ren et al. recently identified several small structured RNAs within the ER lumen, including tRNAs and other RNAs that we previously identified within the glycoRNA pool(*23*). Trafficking through the secretory pathway accompanied by N-glycan trimming and branch extensions would then produce mature sialoglycoRNAs on the cell surface. The mechanisms by which glycoRNAs associate with the plasma membrane are also currently under investigation.

**Fig. 3.**
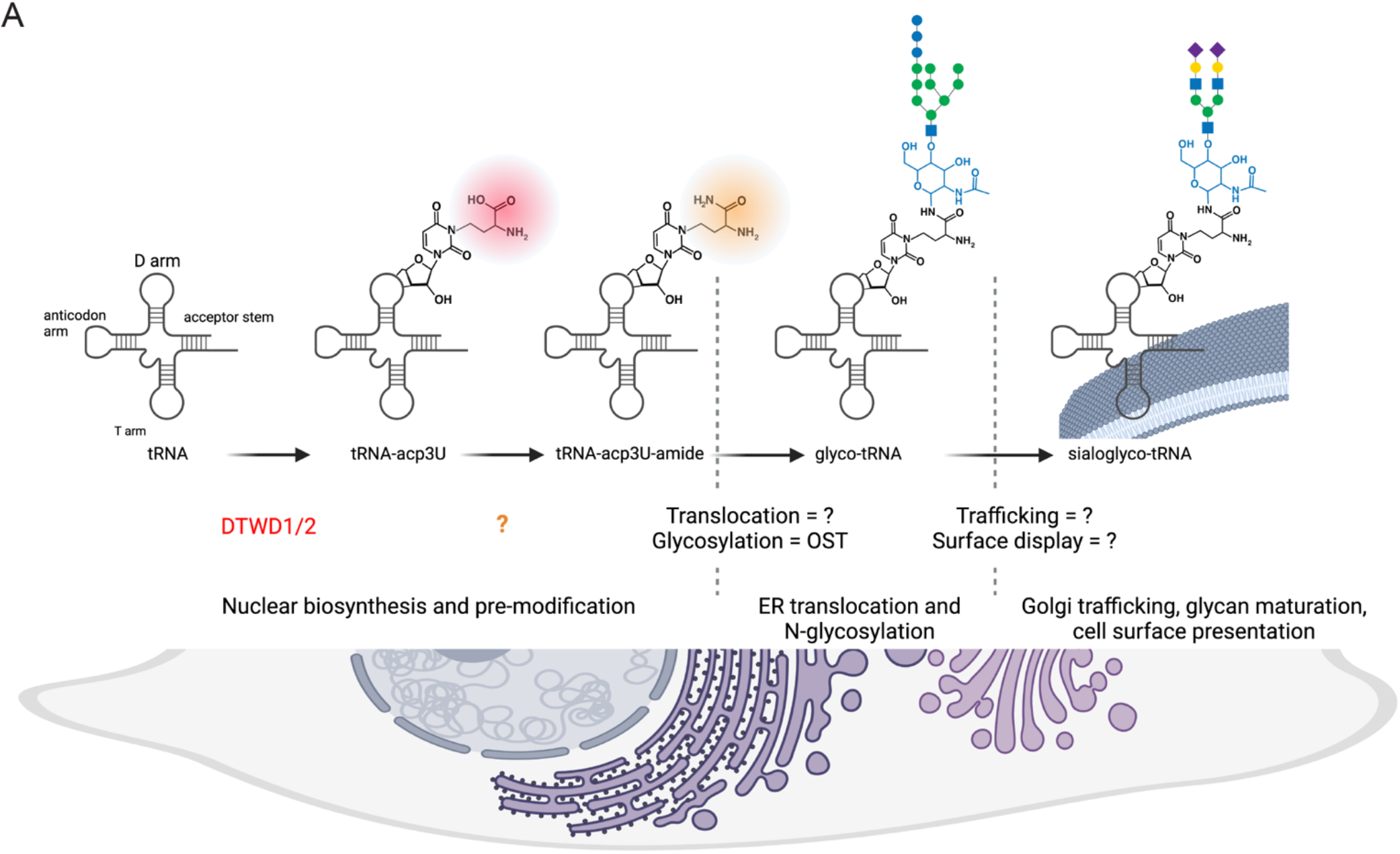
Proposed biosynthetic pathway for generation of cell surface sialoglycoRNAs. (**A**) Cellular tRNAs are synthesized in the nucleus and can be modified there or in the cytosol to create acp3U residue. A subsequent conversion to the carboxamide functionality, by an as-yet unknown enzyme, then allows translocation into the ER lumen. Once in the ER lumen, carboxamide form of acp3U would then enable modification by oligosaccharyltransferase (OST) for N-glycosylation. Further trafficking through the secretory pathway accompanied by N-glycan trimming and branch extensions would then produce mature sialoglycoRNAs on the cell surface.

In proteins, N-glycosylation occurs on asparagine side chains, at the nitrogen atom of the amide functionality. Analogously, we speculate that glucosylation of acp3U would involve prior formation of an amide functionality from the acp3U side chain carboxylic acid. Currently, only one RNA modification, 5-carbamoylmethyluridine (ncm5U), is known to possess a carboxamide group(*12*). However, ncm5U is generated from 5-carboxymethyluridine (cm5U)(*24*), indicating that an amidating enzyme must exist and that other RNA modifications with carboxylic acids may be subject to amidation, making them compatible with N-linked glycosylation. Identification of a putative acp3U amidation pathway is an immediate future goal. Beyond uracil, we found yW-86, a hyper-modified G, as also eluted with PNGaseF (**Fig. 2C, Table S1**) which provides an additional putative site for N-glycans in RNA. yW-86 could offer a linkage mechanism for the putative sites of glycosylation we initially identified via sequencing(*25*).

More broadly, the biosynthesis of sialoglycoRNAs observed in many cell types(*1*, *26*, *27*) suggests that the biochemical activity should be well-conserved. As well, acp3U is found across bacterial and eukaryotic RNAs(*17–19*) which opens the exciting possibility that glycosylated RNAs may exist in non-mammalian species. Despite its conservation, little is known about the cellular role of acp3U: structural analysis suggests it could bind Mg^2+^(*21*) but does not impact the conformation of the ribose(*15*), however more recently mutations in the bacterial acp3U biogenesis (TapT enzyme) cause growth defects in continuous heat stress and double knockouts of DTWD1 and DTWD2 cause growth defects in HEK293T cells(*20*). In this context, we speculate that one of the important cellular roles for acp3U is to provide a site for glycosylation by OST. More broadly, identifying a specific chemical linkage between N-glycans and tRNA brings us one step closer to understanding the biosynthesis and biological functions of glycoRNA.

---

**Table S1. RNA Mass Spectrometry.**

## Acknowledgments

We thank Jonathan Perr, Kayvon Pedram, Stacy Malaker, Maurice Wong, Kevin Janssen, Tiandi Yang, Janelle Sauvageau, and members of the Flynn lab for helpful comments and discussions. We also thank Melissa Gray for the expression and purification of VC-Sialidase. This work was supported by grants from Burroughs Wellcome Fund Career Award for Medical Scientists (R.A.F.), The Rita Allen Foundation (R.A.F.), National Institute of General Medical Sciences of the National Institutes of Health under award number GM151157 (R.A.F.), GM058867 (C.R.B.), AI118891 (B.A.G.), HD106051 (B.A.G.), the Herchel Smith-Harvard Undergraduate Science Research Program (R.M.C.), and an NIH F32 Postdoctoral Fellowship (N.A.T.).

## Author Contributions

R.A.F. conceived the project. R.A.F, B.A.G., and C.R.B. supervised the project and obtained funding. H.H., P.C., C.G.L., R.M.C., and R.A.F. performed RNA labeling optimization experiments and cell based assays. B.M.G. performed PBMC staining and FACS sorting. C.G.L. performed the large scale RNA isolation and labeling. C.P.W. proposed the DTWD pathway and P.C. and C.P.W. developed the knockout cells. C.R.B. and N.A.T. designed the chemical synthesis routes. N.A.T. performed the chemical synthesis, purification, and interpreted NMR and MS data. X.Y. prepared samples for SWATH-MS data. Y.X., B.A.G., R.A.F., N.A.T., and C.R.B. analyzed SWATH-MS data. R.A.F. and C.R.B. wrote the manuscript. All authors discussed the results and revised the manuscript.

## Supplemental Materials

### Supplemental Text

While methods exist to derivatize and label native glycoproteins and glycolipids, no such method exists for the modification of native glycoRNAs. We considered chemoenzymatic labeling, which requires charged sugar donors and specialized enzymes (Reviewed in(*1*)). Instead, we developed an RNA-optimized periodate oxidation and aldehyde labeling technique for the labeling of native glycoRNAs. We establish best practices for handling small RNA samples, develop a one-pot reaction for sialoglycoRNA labeling, and demonstrate its >25-fold increased sensitivity as compared to Ac_4_ManNAz. We further validate that periodate-labeled sialoglycoRNA behaves similarly to our previously reported Ac_4_ManNAz-labeling, establishing this method as an MCR-free alternative to study glycoRNAs.

Over the course of this work, we noticed that precise handling, specific reagents, and optimized procedures dramatically improved reproducibility across our experiments. These features are detailed in the Methods section but major points are noted here. Our previous standard cell lysis used TRIzol, however other reagents like RNAzol RT have been developed to use less phenol and no chloroform in the denaturing and RNA extraction processes(*2*). RNAzol RT extraction of HeLa cells resulted in similar amounts of total RNA isolated per cell compared to TRIzol (**Fig. S1A**). Column clean ups are convenient for rapid processing and are reported to have high RNA recovery. We found that under recommended conditions, HeLa cell total RNA recovery was ∼92% (**Fig. S1B**) while small RNA recovery was < 60% **(Fig. S1C**). We hypothesized this was an issue with the precipitation strength of the prescribed binding reaction and tested various alcohol types and ratios, finding that both ethanol and isopropanol at adjusted ratios could facilitate > 90% recovery of small RNAs using Zymo columns (**Fig. S1C**). Together, these steps represent a series of technical advances that should be broadly applicable to other aspects of RNA biology. With a more reproducible means to isolate and re-purify RNA across chemical reactions, we proceeded to develop a method to label native sialoglycoRNAs.

During the development of this protocol we also re-assessed the need for protease digestion, RNA was extracted without the assistance of proteinase K or StcE during the preparation and only after RNA was isolated, did we then add enzyme to determine co-purification of background proteins and mucins. As we saw previously, proteinase K is important to fully remove proteins from isolated RNA, however while Ac_4_ManNAz signal was not sensitive to mucinase (StcE) digestion(*3*) the PAL signal was (**Fig. S1H**). Additionally, we noticed that our commercially sourced RNA transfer buffer produced incomplete RNA transfers even under the prescribed conditions (**Fig. S2A**). To establish the impact of this, we first screened the pH of the buffer on the transfer properties of Ac_4_ManNAz-labeled RNA. We found that pH < 2 or > 12 led to complete RNA transfer and increased the amount of glycoRNA signal deposited on the membrane (**Fig. S2A**). This increase was apparent but mild and we also noticed additional background bands transferred from Ac_4_ManNAz-labeled RNA at low pH (**Fig. S2A**). We next tested the transfer buffers to verify complete deposition of the rPAL signal onto nitrocellulose membranes. Unlike the ManNAz-labeled signal, which was similarly enhanced at high and low pH, the rPAL signal was slightly weaker using the basic pH buffer conditions while its transfer was dramatically enhanced at low pH (**Fig. S2B**). Further testing defined the optimal pH, salt, and time parameters (**Fig. S2C, S2D**), resulting in an optimized method and enhanced transfer conditions as well as allowing us now to directly compare Ac_4_ManNAz signal to rPAL signal.

These transfer optimization conditions are important as detection of biomolecules with probes or other protein affinity tools fail when the physical transfer from 2D gel separation onto membranes is inefficient. We found that standard methods and buffers to transfer RNA out of denaturing agarose gels were not effective, particularly for rPAL-labeled glycoRNAs. By screening various salt and pH conditions we found a highly efficient process to fully transfer all of the total RNA as well as significantly more sialoglycoRNAs onto the nitrocellulose membrane compared with commercial buffer stocks. Importantly, improved overall transfer of Ac_4_ManNAz-signal resulted in background bands appearing (**Fig. S2A**, left most column) which mirrors some of the background labeling of total RNA we find with rPAL labeling (**Fig. 1C**, “RNA” label). Thus, while neither rPAL or Ac_4_ManNAz is totally background free, the unusual migration of sialoglycoRNAs in the agarose gels enables facile interpretation. Finally, while screening for efficient transfer of Ac_4_ManNAz- vs rPAL-signal, we noticed that Ac_4_ManNAz-signal was enhanced at both high and low pH, while rPAL was transferred poorly at high pH and very efficiently at low pH (**Fig. S2**). This is another difference between Ac_4_ManNAz and rPAL, and suggests some compositional differences in the bulk glycans that are being detected with each method.

We used Ac_4_ManNAz-labeled sialoglycoRNAs as a benchmark, and based on our data, the rPAL-labeled sialoglycoRNAs behave similarly but not identically. rPAL signal fractionated with small RNAs, was completely sialidase sensitive, was reduced in accumulation in cells after inhibiting N-glycosylation biosynthetic enzymes, and importantly rPAL labeled sialoglycoRNAs are highly sensitive to live cell treatment with sialidase, confirming cell surface presentation. However, there are some new features we uncovered using this labeling method. First and most obviously, rPAL labels significantly more sialoglycoRNAs per mass of total RNA from cells, suggesting that Ac_4_ManNAz was relatively inefficient from a fractional incorporation standpoint or rPAL is able to access more sialic acids. The ability to label fractionally more of the sialoglycoRNAs in the cell opened the possibility that we may be detecting novel sialoglycoRNAs in addition to the forms found initially with Ac_4_ManNAz. For example, while we noticed clear and significant loss of rPAL signal after NGI-1 and Kif treatment of HeLa cells, the effects were less than what we had previously seen only with Ac_4_ManNAz-labeling(*3*). Additionally, we previously tested Ac_4_ManNAz-signal for sensitivity to mucinase digestion and saw no effect, while unprocessed RNA labeled with rPAL is partially sensitive to mucinase treatment (**Fig. S1H**). We think therefore that using rPAL can allow researchers to interpret changes in sialoglycoRNA signal to be more directly linked to the biosynthetic flux of sialyltransferases rather than possible differences from cell-type specific variations in metabolic reporter uptake and intracellular usage of Ac_4_ManNAz(*4*). Future work that expands the set of monosaccharides (beyond sialic acid) that can be labeled and investigated in the context of glycoRNA biology will further reveal features of how glycoRNAs operate.

### Methods

#### Cell culture, chemical inhibitor, and metabolic chemical reporters

All cells were grown at 37°C and 5% CO_2_. HeLa cells were cultured in DMEM media supplemented with 10% fetal bovine serum (FBS) and 1% penicillin/streptomycin (P/S). RNA from other cell sources were obtained from^1^ and re-purified as per the descriptions below. Stocks of N-azidoacetylmannosamine-tetraacylated (Ac_4_ManNAz, Click Chemistry Tools) were made to 500 mM in sterile dimethyl sulfoxide (DMSO). For cell treatments, Ac_4_ManNAz was used at a final concentration of 100 μM. Working stocks of glycan-biosynthesis inhibitors were all made in DMSO at the following concentrations and stored at - 80°C: 5 mM NGI-1 (Sigma), 10 mM Kifunensine (Kif, Sigma), 50 mM P-3F_AX_-Neu5Ac (Tocris). All compounds were used on cells for 24 hours.

#### RNA extraction and enzymatic cleanups

TRIzol extractions were performed as previously described in detail(*3*). For RNAzol RT (Molecular Research Center, Inc.) extractions, the manufacturer’s protocol was followed with the following details. First, RNAzol RT was added to lyse and denature cells or tissues, and denaturing was further encouraged by placing the samples at 50°C and shaking for 5 min. To phase separate the RNA, 0.4X volumes of water was added, vortexed, let to stand for 5 minutes at 25°C and lastly spun at 12,000x g at 4°C for 15 min. The aqueous phase was transferred to clean tubes and 1.1X volumes of isopropanol was added. The RNA is then purified over a Zymo column (Zymo Research). We found that preconditioning Zymo columns with water before binding nucleic acids produces more consistent results. For all column cleanups, we followed the following protocol. First, 350 μL of pure water was added to each column and spun at 10,000x g for 30 seconds, and the flowthrough was discarded. Next, precipitated RNA from the RNAzol RT extraction (or binding buffer precipitated RNA, below) is added to the columns, spun at 10,000x g for 10-20 seconds, and the flowthrough is discarded. This step is repeated until all the precipitated RNA is passed over the column once. Next, the column is washed three times total: once using 400 μL RNA Prep Buffer (3M GuHCl in 80% EtOH), twice with 400 μL 80% ethanol. The first two spins are at 10,000x g for 20 seconds, the last for 30 sec. The RNA is then treated with Proteinase K (Ambion) on the column. Proteinase K is diluted 1:19 in water and added directly to the column matrix (Zymo-I = 20 μL, Zymo-II = 50 μL, Zymo-IIICG = 60 μL, all from Zymo Research), and then allowed to incubate on the column at 37°C for 45 min. The column top is sealed with either a cap or parafilm to avoid evaporation. After the digestion, the columns are brought to room temperature for 5 min; lowering the temperature is important before proceeding. Next, eluted RNA is spun out into fresh tubes and a second elution with water is performed (Zymo-I = 30 μL, Zymo-II = 50 μL, Zymo-IIICG = 60 μL). To the eluate, 1.5 μg of the mucinase StcE (Sigma-Aldrich) is added for every 50 μL of RNA, and placed at 37°C for 30 minutes to digest. The RNA is then cleaned up again using a Zymo column. Here, 2X RNA Binding Buffer (Zymo Research) was added and vortexed for 10 seconds, and then 2X (samples + buffer) of 100% ethanol was added and vortexed for 10 sec. An example would be 50 μL of RNA, 100 μL of RNA Binding Buffer, and 300 μL of 100% ethanol. This is then bound to the column, cleaned up as described above, and eluted twice with water (Zymo-I = 25 μL, Zymo-II = 50 μL, Zymo-IIICG = 60 μL). The final enzymatically digested RNA is quantified using a Nanodrop.

Binding conditions to efficiently precipitate small RNAs as highlighted in **Fig. S1C** were optimized by varying the amount of ethanol added post RNA Binding Buffer mixing with RNA. Isopropanol was also exchanged for the ethanol at this step to assess its ability to facilitate small RNA capture on the Zymo columns.

After total RNA extraction, the RNA can be further processed in order to fractionate small (17-200 nts) and large RNA (>200 nts) using Zymo columns. First, an adjusted RNA binding buffer is made by mixing equal volumes of RNA binding buffer and 100% ethanol. Two volumes of the adjusted buffer are added to the total RNA and are vortexed thoroughly to mix. The sample is then bound to the column as described above, but the flow through - which contains the small RNA - is saved. One volume of 100% ethanol is added to the small RNA and vortexed to mix. The small RNA is then bound to new columns. Both the large and small RNA (bound to their respective columns) are then cleaned up using 2X 400 μL 80% ethanol, the second clean being centrifuged for 30 seconds. The RNA is then eluted using 2X 50 μL of water.

#### In vitro and On Cell enzymatic digestions

To digest RNA, the following was used: 2 μL of RNase cocktail (0.5U/mL RNaseA and 20U/mL RNase T1, Thermo Fisher Scientific) with 20 mM Tris-HCl (pH 8.0), 100 mM KCl and 0.1 mM MgCl_2_. To digest sialic acid: 1.5 μL of a2-3,6,8 Neuraminidase (50U/mL, New England Biolabs, NEB) with 1x GlycoBuffer 1 (NEB). Reactions were performed on 3 μg total RNA from indicated cell sources for 60 minutes at 37°C. For live cell treatments, VC-Sia was expressed and purified as previously described(*5*) and added to cells at 150 nM final concentration in complete growth media for between 15 and 60 minutes at 37°C.

#### Isolation of crude cellular membranes

Crude membranes were isolated using the Plasma Membrane Protein Extraction Kit (ab65400, Abcam): cultured cells first had growth media removed and cells were then washed twice with ice-cold 1x Phosphate Buffered Saline (PBS). In the second PBS wash, cells were scraped off the plate and spun down at 400x g for 4 minutes at 4°C or suspension cells were directly pelleted. Cell pellets were resuspended in 1mL of Homogenization Buffer Mix per 10,000,000 cells. Cell suspension was Dounce Homogenized on ice for 40-70 strokes, care was taken to stop douncing when the processing resulted in approximately 60% free nuclei so as to not generate excess lysis of nuclei. Homogenate was then spun at 700x g for 10 minutes at 4°C. This pellet contained the nuclear fraction and supernatants were transferred to new tubes and spun again at 10,000x g for 30 minutes at 4°C. The pellets generated from this spin were crude membranes and the supernatant was soluble cytosol. RNA extraction was performed as above and labeling as described below.

#### Periodate oxidation and aldehyde ligation for glycoRNA labeling

Starting with a maximum of 3 ug of lyophilized, enzymatically treated RNA, the first step of the rPAL labeling protocol is to block any aldehyde reactive species. To make the blocking buffer, 1μL 16 mM mPEG3-Ald (BP-23750, BroadPharm), 15 μL 1 M MgSO_4_ and 12 μL 1 M NH_4_OAc pH5 (with HCl) are mixed together (final buffer composition: 570 μM mPEG3-Ald + 500 mM MgSO_4_ + 450 mM NH_4_OAc pH5). 28μL of the blocking buffer is added to the lyophilized RNA, mixed completely by vortexing, and then incubated for 45 minutes at 35°C to block. The samples are briefly allowed to cool to room temperature (2-3 min), then working quickly, 1 μL 30 mM aldehyde reactive probe (ARP/aminooxy biotin, 10009350, Cayman Chemicals, stock made in water) is added first, then 2μL of 7.5 mM NaIO_4_ (periodate, stock made in water) is added. The periodate is allowed to perform oxidation for exactly 10 minutes at room temperature in the dark. The periodate is then quenched by adding 3 μL of 22 mM sodium sulfite (stock made in water). The quenching reaction is allowed to proceed for 5 minutes at 25°C. Both the sodium periodate and sodium sulfite stocks were made fresh within 20 minutes of use. Next, the reactions are moved back to the 35°C heat block, and the ligation reaction is allowed to occur for 90 min. The reaction is then cleaned up using a Zymo-I column. 19 μL of water is added in order to bring the reaction volume to 50 μL, and the Zymo protocol is followed as per the above details. If samples will be analyzed on an agarose gel for glycoRNA visualization, the RNA is then eluted from the column using 2X 6.2μL water (final volume approximately 12 μL).

#### glycoRNA blotting

In order to visualize the periodate labeled RNA, it is run on a denaturing agarose gel, transferred to a nitrocellulose (NC) membrane, and stained with streptavidin in a manner similar to (*3*) with some modifications. After elution from the column as described above, the RNA is combined with 12 μL of Gel Loading Buffer II (GLBII, 95% formamide, 18 mM EDTA, 0.025% SDS) with a final concentration of 2X SybrGold (ThermoFisher Scientific) and denatured at 55°C for 10 minutes. It is important to not use GLBII with dyes. Immediately after this incubation, the RNA is placed on ice for at least 2 minutes. The samples are then loaded into a 1% agarose, 0.75% formaldehyde, 1.5x MOPS Buffer (Lonza) denaturing gel. Precise and consistent pouring of these gels is critical to ensure a similar thickness of the gel for accurate transfer conditions; we aim for approximately 1 cm thick of solidified gel. RNA is electrophoresed in 1x MOPS at 115V for between 34 or 45 min, depending on the length of the gel. Subsequently, the RNA is visualized on a UV gel imager, and excess gel is cut away; leaving ∼0.75 cm of gel around the outer edges of sample lanes will improve transfer accuracy. The RNA is transferred as previously described(*3*), however various buffer conditions and times were screened to determine the optimal method (**Fig. S2**). Finally, we determined that a 3M NaCl solution at pH 1, achieved with HCl, yields the most consistent and efficient transfer of material to the NC membrane. Transfer occurs for 90 minutes at 25°C. Post transfer, the membrane is rinsed in 1x PBS and dried on Whatman Paper (GE Healthcare). Dried membranes are rehydrated in Intercept Protein-Free Blocking Buffer, TBS (Li-Cor Biosciences), for 30 minutes at 25°C. After the blocking, the membranes are stained using Streptavidin-IR800 (Li-Cor Biosciences) diluted 1:5,000 in Intercept blocking buffer for 30 minutes at 25°C. Excess Streptavidin-IR800 was washed from the membranes using three washes with 0.1% Tween-20 (Sigma) in 1x PBS for 3 minutes each at 25°C. The membranes were then briefly rinsed with PBS to remove the Tween-20 before scanning. Membranes were scanned on a Li-Cor Odyssey CLx scanner (Li-Cor Biosciences).

#### Peripheral blood mononuclear cell staining and flow cytometry sorting

Human Peripheral Blood Mononuclear Cells (PBMCs) were purchased from StemCell Technology. Samples from three individual donors were thawed at 37°C for 3 minutes and then diluted into 5 mL of FACS buffer (0.5% BSA in 1x PBS). Freezing media was removed by spinning the cells for 4 minutes at 400g at 4°C. 10 million cells were resuspended in 1 mL of FACS buffer and 50 μL of Human TruStain FcX Fc Receptor Blocking Solution (BioLegend) was added and left to bind the cells on ice for 15 minutes. The blocked PMBCs were stained with anti-CD19 clone HIB19 conjugated to PE-Cy7 (BioLegend), anti-CD3 clone OKT3 conjugated to FITC (BioLegend), anti-CD14 clone HCD14 conjugated to APC-Cy7 (BioLegend), and anti-CD16 clone 3G8 conjugated to PerCP-Cy5.5 (BioLegend) for 30 minutes on ice. Cells were pelleted by spinning for 4 minutes at 400g at 4°C. The supernatant was discarded and the cells were resuspended in 1 mL of FACS buffer supplemented with 4’,6-Diamidino-2-phenylindole (DAPI) at 1 μg/mL final concentration as a live/dead stain. Stained cells were sorted into four major subpopulations using a Sony MA900 sorter with a 100 μM sorting chip (Sony). Cells were collected into FACS buffer and spun for 4 minutes at 400g at 4°C. Pellets were lysed in RNAzol RT and rPAL labeling was performed as described above.

#### rPAL labeling, glycoRNA capture, and enzymatic release

RNA from HEK293 and K562 cells was isolated, enzymatically digested, and removed of large RNA as described above. For each replicate 250 μg of small RNA was used. The rPAL procedure was performed as described above, scaled up linearly, with a 1x reaction assuming 3 μg input RNA. We performed the rPAL with or without the aminooxy-biotin reagent, omitting this reagent served as a control sample. After the rPAL labeling and Zymo cleanups were accomplished we captured biotinylated RNAs using 250 μL Neutravidin bead slurry (ThermoFisher Scientific) per replicate. The capture took place at 4°C over 2 hours on rotation in a final volume of 1000 μL in 5x PBS. Beads were then washed sequentially with 1000 μL of the following buffers for 30 seconds each, all pre-warmed to 45°C before use: twice with 4 M NaCl, 100 mM HEPES, then twice 2 M GuHCl, 100 mM HEPES, then twice with room temperature 1x PBS. The washed beads were then subjected to nuclease treatment by incubating in 300 μL of 250 μg RNase A (Sigma, 10109169001), 5 units of Phosphodiesterase (PDE1, Sigma, P3243-1VL), and 0.5 mM MgCl_2_ in 0.5x PBS. Nuclease digestion occurred at 37°C for 4 hours with shaking to 1100 rpm for 10 sections, every 60 sections. The supernatant of this reaction was collected in a new tube and the beads washed with 1000 μL of LC-MS grade water (ThermoFisher Scientific) and the water supernatant of this wash was pooled with the nuclease released supernatant. The pooled supernatants were frozen and lyophilized for LC-MS analysis as described below. The nuclease digested beads were next subjected to one of two types of glycosidase treatments, either PNGaseF or Sialidase, both of which occurred at 37°C for 16 hours with shaking to 1100 rpm for 10 sections, every 60 sections. Sialidase digestion mix was a final volume of 250 μL in 1x NEB Glyco Buffer #1 with 5 μL of α2-3,6,8 Neuraminidase (NEB, P0720) and 5 μL of α2-3,6,8,9 Neuraminidase A (NEB, P0722). PNGaseF digestion mix was a final volume of 250 μL in 1x PBS with 7.5 μL PNGaseF prime (Bulldog Bio, NZPP050LY). After the overnight reaction, the supernatants were collected again in a fresh tube, beads washed with LC-MS water as above, supernatants pooled, and samples lyophilized for LC-MS analysis.

#### Porous Graphitic Carbon Liquid Chromatography Mass Spectrometry Analysis

##### Nucleosides sample preparation

10 mU of shrimp alkaline phosphatase (SAP, New England Biolabs) was added to the samples generated above to remove the phosphates. The reaction was employed in 50 mM sodium acetate (pH 7.2) for 2 hours at 37 °C. The nucleoside samples were further purified using the Hypercarb SPE 96-well plate (Thermo Fisher Scientific). The plate was first activated and equilibrated with a solution of 80% (v/v) Acetonitrile (ACN, LC-MS Grade, Thermo Fisher Scientific) and 0.1% (v/v) TFA and 0.1% (v/v) Trifluoroacetic Acid (TFA, LC-MS Grade, Thermo Fisher Scientific) in LC-MS water, respectively. The dried samples were solubilized, loaded onto the cartridge, and washed with 0.1% (v/v) TFA in water. Nucleosides were eluted with a solution of 60% (v/v) ACN and 0.1% (v/v) TFA and dried using the TurboVap LV Evaporator (Biotage, Sweden).

##### LC-MS/MS Analysis

The digested nucleosides were reconstituted in MS buffer A (LC-MS water with 0.1% Formic Acid (FA, LC-MS Grade, Fisher Scientific) (v/v)), and 2 μL was injected for the analysis using the SWAMNA platform(*6*). Briefly, the samples were introduced using nanoAcquity UPLC System (Waters) equipped with Hypercarb Porous Graphitic Carbon HPLC Column (1 mm x 100 mm, 3 μm). Separation was carried out at a constant flow rate of 40 μL/min using MS buffer A and buffer B (ACN with 0.1% FA (v/v). The gradient was referred from the previous protocol<u>^21^</u>. Specifically, 0−2 min, 0% B; 2−20 min, 0-16% B; 20−40 min, 16%-72% B; 40−42 min, 72%-100% B; 42-52 min, 100% B; 52-54 min 100%-0% B; 54-65 min 0% B. The analytes were ionized using OptiFlow Turbo V ion source (SCIEX) at 5000 V and analyzed using ZenoTOF 7600 (SCIEX). Ion source gas1, gas2, and curtain gas were at 40, 60, and 35 psi, respectively. The ion transfer tube temperature was set at 200 °C. For zenoSWATH analysis, MS spectra were acquired over a mass range of m/z 200–1250 in positive ionization mode with 0.25 s accumulation time. Collision-induced fragmentation was performed with nitrogen gas using dynamic collision energy. The product ions were monitored from the range m/z 100-1200 with 15 ms accumulation time. For the targeted MRMHR analysis, the monitored masses and chargers were listed in **Table S1**.

#### Data Analysis

The data was manually inspected using SCIEX OS Explorer (v3.0), NuMo Finder, and Skyline (v22.2)(*7*). Add raw data has been deposited in the MassIVE database with accession number: MSV000093213.

##### Generation of KO U2OS cell lines

DTWD2 KO cell lines were generated using a CRISPR/Cas9 approach(*8*). The target oligonucleotide (5’-AGCGCACCCTCTGCATATCT-3’) was synthesized and ligated into PX459 vectors. U2OS cells were then transfected with gRNA vectors. Two days later, puromycin (2 µg/mL) was added into the cell culture and the live cells were selected by flow cytometry (FACS Calibur 2, BD science) for isolation of single clones. The expanded individual clones were screened by genomic DNA sequencing.

### General Information for Synthetic Procedures

Peracetylated azido-GlcNAc (2-Acetamido-3,4,6-tri-O-acetyl-2-deoxy-β-D-glucopyranosyl azide) and Boc-homoserine-CO_2_Me ((*S*)-Methyl 2-((tert-butoxycarbonyl)amino)-4-hydroxybutanoate) were purchased from Synthose (catalog # AG931) and Chem Scene (catalog # CS-0088997) respectively and used without further purification. Acetylated amino-GlcNAc ((2*R*,3*S*,5*R*,6*R*)-5-acetamido-2-(acetoxymethyl)-6-aminotetrahydro-2H-pyran-3,4-diyl diacetate, **S6**) was prepared by palladium-catalyzed hydrogenation of the corresponding commercially available azide starting material according to a literature procedure(*9*). Solvents were purified with a PureSolv Solvent Purification System or used as commercially available anhydrous sure-seal bottles. Non-aqueous reagents were transferred under nitrogen via syringe. Organic solutions were concentrated under reduced pressure on a Büchi rotary evaporator using a water bath. Normal phase chromatographic purification of products was accomplished using forced-flow chromatography on a Biotage® automated flash column chromatography system equipped with Biotage® Sfär pre-packed silica gel columns (20 μm particle size). Thin-layer chromatography (TLC) was performed on Silicycle 0.25 mm silica gel F-254 plates. Visualization of the developed chromatogram was performed by UV lamp exposure and KMnO_4_ stain. ^1^H NMR spectra were recorded on a Bruker NEO-500 MHz and are internally referenced to residual protio D_2_O (4.79 ppm), CD_3_OD (3.31 ppm), or CDCl_3_ (7.26 ppm) signals. CDCl_3_ was stored over K_2_CO_3_. Data for ^1^H NMR are reported as follows: chemical shift (δ ppm), integration, multiplicity (s = singlet, d = doublet, t = triplet, q = quartet, m = multiplet, dd = doublet of doublets, dt = doublet of triplets, br = broad), and coupling constant (Hz). ^13^C NMR spectra were recorded on a Bruker UltraShield Plus 500 MHz and data are reported in terms of chemical shift relative to CDCl_3_ (77.16 ppm) or CD_3_OD (49.00 ppm). HRMS (High Resolution Mass Spectrometry) data were obtained on an Agilent 6230B Time of Flight (TOF) LC/MS by direct infusion.

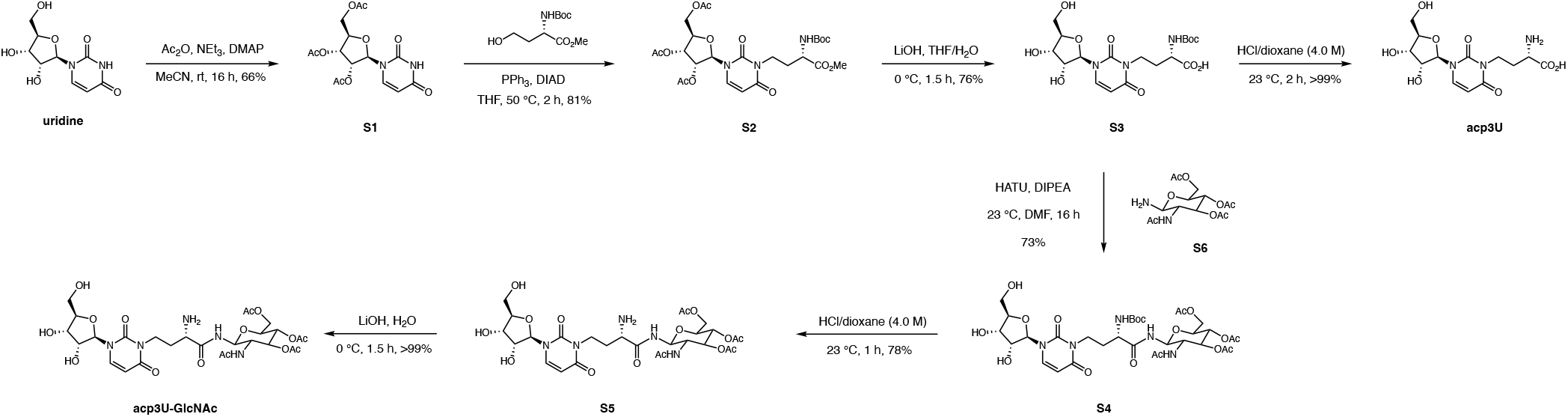
Overview of synthetic approach to acp3U and acp3U-GlcNAc.

### Synthetic Procedures and Characterization Data

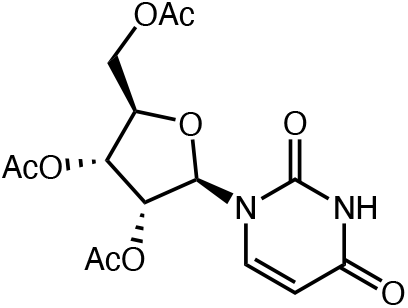

### (2*R*,3*R*,4*R*,5*R*)-2-(acetoxymethyl)-5-(2,4-dioxo-3,4-dihydropyrimidin-1(2H)-yl)tetrahydrofuran-3,4-diyl diacetate (S1)

To a round bottom flask charged with starting uridine (5.00 g, 20.5 mmol), 40 mL anhydrous acetonitrile was added followed by triethylamine (4.00 equiv, 11.4 mL, 81.9 mmol), acetic anhydride (3.00 equiv, 7.63 mL, 61.4 mmol), and 4-dimethylaminopyridine (DMAP, 1 mol%, 25 mg, 205 μmol) in that order. The reaction was left to stir at room temperature (23 °C) for 16 hours before concentrating and redissolving in chloroform (100 mL). The organic layer was washed with water (2 X 50 mL), dried over MgSO_4_, and purified by SiO_2_ gel column chromatography (0–3% MeOH/DCM) to yield a dark brown syrup. This material was subjected to a second identical SiO_2_ gel column to yield a beige solid (5.0 g, 66% yield). ^1^H NMR (500 MHz, CDCl3) δ 8.56 (s, 1H), 7.39 (d, J = 8.2 Hz, 1H), 6.06 – 6.01 (m, 1H), 5.79 (dd, J = 8.1, 1.5 Hz, 1H), 5.36 – 5.31 (m, 2H), 4.41 – 4.31 (m, 3H), 2.15 (s, 3H), 2.13 (s, 3H), 2.10 (s, 3H); ^13^C NMR (126 MHz, CDCl3) δ 170.26, 169.78, 169.77, 162.77, 150.29, 139.40, 103.57, 87.58, 80.08, 72.85, 70.32, 63.28, 20.91, 20.64, 20.55; HRMS (ESI-TOF) *m*/*z* calcd. for C_15_H_18_N_2_O_9_Na ([M+Na]^+^) 393.091003, found 393.0893.

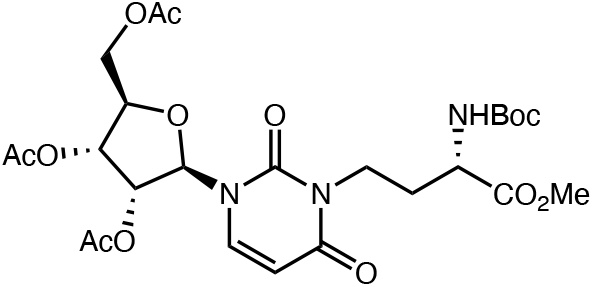

### (2*R*,3*R*,4*R*,5*R*)-2-(acetoxymethyl)-5-(3-((*S*)-3-((tert-butoxycarbonyl)amino)-4-methoxy-4-oxobutyl)-2,4-dioxo-3,4-dihydropyrimidin-1(2H)-yl)tetrahydrofuran-3,4-diyl diacetate (S2)

To a round bottom flask charged with peracetylated uridine **S1** (661 mg, 1.79 mmol), 50 mL anhydrous THF was added, followed by Boc-homoserine-CO_2_Me ((*S*)-Methyl 2-((tert-butoxycarbonyl)amino)-4-hydroxybutanoate) (1.2 equiv, 500 mg, 2.14 mmol), triphenylphosphine (1.2 equiv, 562 mg, 2.14 mmol), and DIAD (diisopropyl azodicarboxylate) (1.2 equiv, 421 μL, 2.14 mmol) in that order. The solution was stirred at 50 °C for 2 hours before concentration, and subjection to SiO_2_ gel column chromatography (0– 50% acetone/hexane) to give a white solid contaminated with 50 wt% PPh_3_O triphenylphosphine oxide (842 mg desired product, 81% yield). This mixture was carried forward without further purification. ^1^H NMR (500 MHz, CDCl3) δ 7.36 (d, J = 8.1 Hz, 1H), 6.03 (d, J = 5.1 Hz, 1H), 5.79 (d, J = 8.2 Hz, 1H), 5.45 (d, J = 8.8 Hz, 1H), 5.37 (t, J = 5.5 Hz, 1H), 5.34 – 5.30 (m, 1H), 4.44 – 4.32 (m, 4H), 4.13 – 4.03 (m, 1H), 4.00 – 3.92 (m, 1H), 3.67 (s, 3H), 2.17 – 2.09 (m, 11H), 1.45 (s, 9H); ^13^C NMR (126 MHz, CDCl3) δ 172.71, 170.22, 169.76, 169.73, 162.21, 155.69, 150.89, 137.42, 102.79, 88.55, 80.06, 79.96, 72.97, 70.17, 63.13, 52.49, 51.46, 37.75, 29.62, 28.47, 20.92, 20.65, 20.60; HRMS (ESI-TOF) *m*/*z* calcd. For C_25_H_35_N_3_O_13_Na add([M+Na]^+^) 608.206762, found 608.2094.

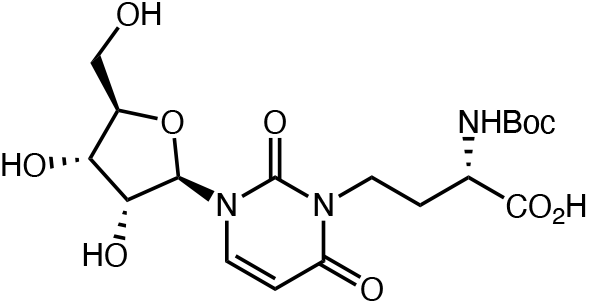

### (*S*)-2-((tert-butoxycarbonyl)amino)-4-(3-((2*R*,3*R*,4*S*,5*R*)-3,4-dihydroxy-5-(hydroxymethyl)tetrahydrofuran-2-yl)-2,6-dioxo-3,6-dihydropyrimidin-1(2H)-yl)butanoic acid (S3)

Starting Boc-acp3U methyl ester **S2** (250 mg, 427 μmol) was dissolved in 4.30 mL THF and cooled on ice to 0 °C. To this solution, LiOH was added as an aqueous 1.00 M solution (5.00 equiv, 2.13 mmol) at 0 °C. The solution was stirred at 0 °C for 90 minutes before neutralizing to pH 7 by the addition of 10% aq. HCl. THF was removed from the solution by rotary evaporation before the aqueous mixture was purified by preparative scale reversed-phase HPLC (C18, 0-20% MeCN/0.1%FA in H_2_O/0.1%FA). Product-containing fractions were combined, MeCN was removed by rotary evaporation, and lyophilized to give the title compound as a white solid (145 mg, 76% yield). ^1^H NMR (500 MHz, D_2_O) δ 7.87 (d, J = 8.1 Hz, 1H), 5.94 (d, J = 8.1 Hz, 1H), 5.91 (d, J = 3.9 Hz, 1H), 4.35 (dd, J = 5.3, 4.0 Hz, 1H), 4.22 (t, J = 5.7 Hz, 1H), 4.16 – 4.12 (m, 2H), 4.06 (t, J = 6.9 Hz, 2H), 3.94 (dd, J = 12.8, 2.9 Hz, 1H), 3.82 (dd, J = 12.8, 4.4 Hz, 1H), 2.28 – 2.15 (m, 1H), 2.12 – 2.02 (m, 1H), 1.45 (s, 9H); ^13^C NMR (126 MHz, D_2_O) δ 176.28, 165.08, 157.55, 151.67, 139.81, 101.58, 90.46, 83.93, 81.46, 73.72, 69.19, 60.58, 52.09, 38.19, 29.58, 28.11, 27.61; HRMS (ESI-TOF) *m*/*z* calcd. for C_18_H_27_N_3_O_10_Na ([M+Na]^+^) 468.159416, found 468.1574.

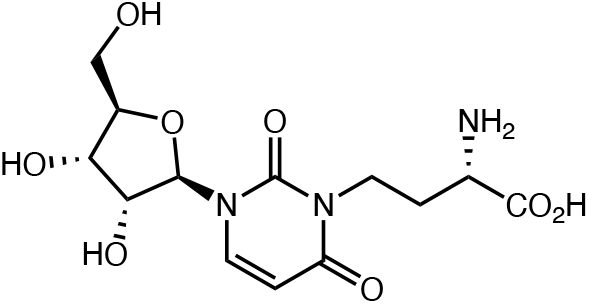

### (*S*)-2-amino-4-(3-((2*R*,3*R*,4*S*,5*R*)-3,4-dihydroxy-5-(hydroxymethyl)tetrahydrofuran-2-yl)-2,6-dioxo-3,6-dihydropyrimidin-1(2H)-yl)butanoic acid (acp3U)

To a 4 mL vial charged with Boc-protected acp3U **S3** (20.5 mg, 46.0 μmol), 2.0 mL 4.0 M HCl/dioxane was added. The solution was stirred at room temperature (23 °C) for 2 hours before concentration. The resulting residue was redissolved in H_2_O and purified by preparative scale reversed-phase HPLC (C18, 0-20% MeCN/0.1%FA in H_2_O/0.1%FA). Product-containing fractions were combined, MeCN was removed by rotary evaporation, and lyophilized to give the title compound as a white solid (15.9 mg, >99% yield). ^1^H NMR (500 MHz, D_2_O) δ 7.89 (d, J = 8.1 Hz, 1H), 5.97 (d, J = 8.1 Hz, 1H), 5.93 (d, J = 4.0 Hz, 1H), 4.37 (dd, J = 5.4, 4.1 Hz, 1H), 4.23 (t, J = 5.6 Hz, 1H), 4.17 – 4.04 (m, 3H), 3.94 (dd, J = 12.8, 2.8 Hz, 1H), 3.85 – 3.79 (m, 2H), 2.34 – 2.20 (m, 2H); ^13^C NMR (126 MHz, D_2_O) δ 173.28, 165.23, 151.84, 139.96, 101.59, 90.35, 84.03, 73.73, 69.22, 60.58, 52.13, 37.36, 27.79; HRMS (ESI-TOF) *m*/*z* calcd. for C_13_H_20_N_3_O_8_ ([M+H]^+^) 346.125042, found 346.1243.

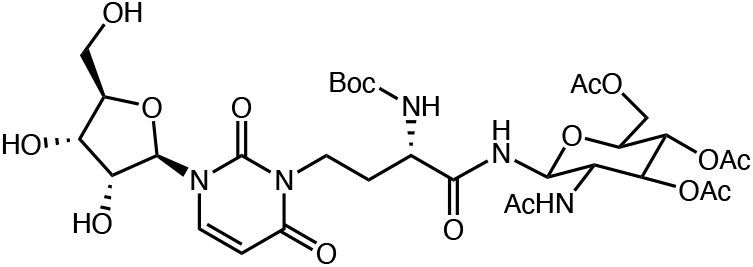

### (2*S*,3*R*,5*S*,6*S*)-5-acetamido-2-(acetoxymethyl)-6-((*S*)-2-((tert-butoxycarbonyl)amino)-4-(3-((2*R*,3*R*,4*S*,5*R*)-3,4-dihydroxy-5-(hydroxymethyl)tetrahydrofuran-2-yl)-2,6-dioxo-3,6-dihydropyrimidin-1(2H)-yl)butanamido)tetrahydro-2H-pyran-3,4-diyl diacetate (S4)

A 20 mL scintillation vial was charged with starting Boc-acp3U **S3** (50.0 mg, 112 μmol), peracetylated GlcNAc-NH_2_ **S6** (1.20 equiv, 46.7 mg, 135 μmol), and HATU (1.00 equiv, 42.7 mg, 135 μmol). To this vial 2 mL DMF was added followed by DIPEA (4.00 equiv, 78.2 μL, 449 μmol). The mixture was stirred at room temperature (23 °C) for 16 hours before direct subjection to preparative scale reversed-phase HPLC purification (C18, 0-50% MeCN/0.1%FA in H_2_O/0.1%FA). Product-containing fractions were combined, MeCN was removed by rotary evaporation, and lyophilized to give the title compound as a white solid (63.1 mg, 73% yield). ^1^H NMR (500 MHz, CD_3_OD) δ 8.05 (d, J = 8.1 Hz, 1H), 5.91 (d, J = 4.0 Hz, 1H), 5.77 (d, J = 8.2 Hz, 1H), 5.23 – 5.17 (m, 2H), 5.00 (d, J = 9.3 Hz, 1H), 4.29 – 4.23 (m, 1H), 4.19 (t, J = 4.6 Hz, 1H), 4.15 (t, J = 5.3 Hz, 1H), 4.11 – 4.05 (m, 2H), 4.04 – 3.94 (m, 4H), 3.89 – 3.81 (m, 2H), 3.75 (dd, J = 12.2, 3.1 Hz, 1H), 2.05 – 1.89 (m, 14H), 1.44 (s, 9H); ^13^C NMR (126 MHz, MeOD) δ 175.26, 174.04, 172.37, 171.82, 171.31, 164.92, 157.60, 152.70, 140.86, 102.01, 91.76, 86.16, 80.66, 79.89, 75.90, 74.74, 74.68, 70.90, 69.95, 63.24, 61.95, 54.34, 53.98, 39.02, 30.89, 28.73, 22.78, 20.64, 20.58, 20.55. HRMS (ESI-TOF) *m*/*z* calcd. for C_32_H_47_N_5_O_17_Na ([M+Na]^+^) 796.28647, found 796.2886.

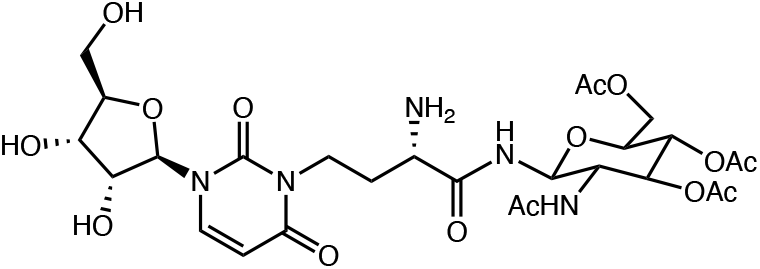

### (2*S*,3*R*,5*S*,6*S*)-5-acetamido-2-(acetoxymethyl)-6-((*S*)-2-amino-4-(3-((2*R*,3*R*,4*S*,5*R*)-3,4-dihydroxy-5-(hydroxymethyl)tetrahydrofuran-2-yl)-2,6-dioxo-3,6-dihydropyrimidin-1(2H)-yl)butanamido)tetrahydro-2H-pyran-3,4-diyl diacetate (S5)

To a 4 mL vial charged with Boc-protected starting material **S4** (9.9 mg, 12.8 μmol), 2.0 mL 4.0 M HCl/dioxane was added. The solution was stirred at room temperature (23 °C) for 60 minutes before concentration. The resulting residue was redissolved in H_2_O and purified by preparative scale reversed-phase HPLC (C18, 0-20% MeCN/0.1%FA in H_2_O/0.1%FA). Product-containing fractions were combined, MeCN was removed by rotary evaporation, and lyophilized to give the formate salt of the title compound as a white solid (7.2 mg, 78% yield). ^1^H NMR (500 MHz, D_2_O) δ 7.91 (d, J = 8.1 Hz, 1H), 5.98 (d, J = 8.1 Hz, 1H), 5.95 (d, J = 4.3 Hz, 1H), 5.33 (t, 2H), 5.11 (t, J = 9.7 Hz, 1H), 4.40 – 4.35 (m, 2H), 4.25 – 4.02 (m, 7H), 3.98 – 3.91 (m, 2H), 3.83 (dd, J = 12.8, 4.3 Hz, 1H), 2.32 – 2.15 (m, 2H), 2.13 (s, 3H), 2.11 (s, 3H), 2.08 (s, 3H), 1.96 (s, 3H); ^13^C NMR (126 MHz, D_2_O) δ 174.59, 173.69, 173.12, 172.83, 171.00, 165.13, 151.89, 139.98, 101.65, 90.06, 84.14, 77.94, 73.76, 73.43, 72.95, 69.26, 68.28, 61.93, 60.56, 52.09, 51.29, 37.16, 28.92, 21.86, 20.11, 20.06, 19.96; HRMS (ESI-TOF) *m*/*z* calcd. for C_27_H_40_N_5_O_15_ ([M+H]^+^) 674.252095, found 674.2539.

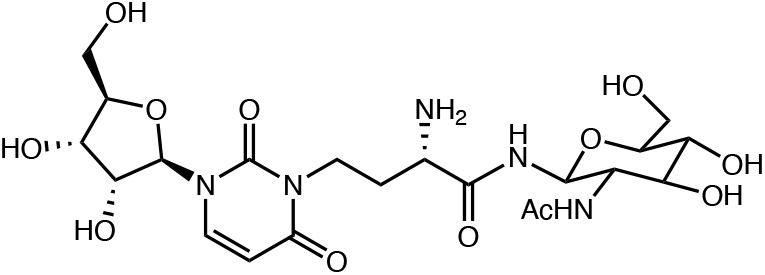

### (2*S*)-N-((2*S*,3*S*,5*R*,6*S*)-3-acetamido-4,5-dihydroxy-6-(hydroxymethyl)tetrahydro-2H-pyran-2-yl)-2-amino-4-(3-((2R,3R,4S,5R)-3,4-dihydroxy-5-(hydroxymethyl)tetrahydrofuran-2-yl)-2,6-dioxo-3,6-dihydropyrimidin-1(2H)-yl)butanamide (acp3U-GlcNAc)

Starting acetylated acp3U-GlcNAc **S5** (9.3 mg, 427 μmol) was dissolved in 2.0 mL H_2_O and cooled on ice to 0 °C. To this solution, LiOH was added as an aqueous 1.00 M solution (5.00 equiv, 69.0 μmol) at 0 °C. The solution was stirred at 0 °C for 90 minutes before neutralizing to pH 7 by the addition of 10% aq. HCl. The neutralized mixture was purified by preparative scale reversed-phase HPLC (C18, 0-20% MeCN/0.1%FA in H_2_O/0.1%FA). Product-containing fractions were combined, MeCN was removed by rotary evaporation, and lyophilized to give the formate salt of the title compound as a white solid (8.4 mg, >99% yield). ^1^H NMR (500 MHz, D_2_O) δ 7.91 (d, J = 8.2 Hz, 1H), 5.99 (d, J = 8.1 Hz, 1H), 5.95 (d, J = 4.1 Hz, 1H), 5.09 (d, J = 9.8 Hz, 1H), 4.37 (t, J = 4.8 Hz, 1H), 4.23 (t, J = 5.5 Hz, 1H), 4.18 – 4.13 (m, 1H), 4.13 – 4.02 (m, 2H), 3.97 – 3.86 (m, 4H), 3.83 (dd, J = 12.8, 4.4 Hz, 1H), 3.77 (dd, J = 12.3, 4.4 Hz, 1H), 3.67 – 3.60 (m, 1H), 3.56 – 3.48 (m, 2H), 2.34 – 2.10 (m, 2H), 2.01 (s, 3H); ^13^C NMR (126 MHz, D_2_O) δ 174.84, 171.00, 165.18, 151.90, 140.02, 101.67, 90.14, 84.12, 78.58, 77.68, 74.19, 73.76, 69.39, 69.25, 60.58, 60.46, 54.13, 51.29, 37.15, 28.83, 22.06; HRMS (ESI-TOF) *m*/*z* calcd. for C_21_H_33_N_5_O_12_Na ([M+Na]^+^) 570.202344, found 570.2046.

**Fig. S1.**
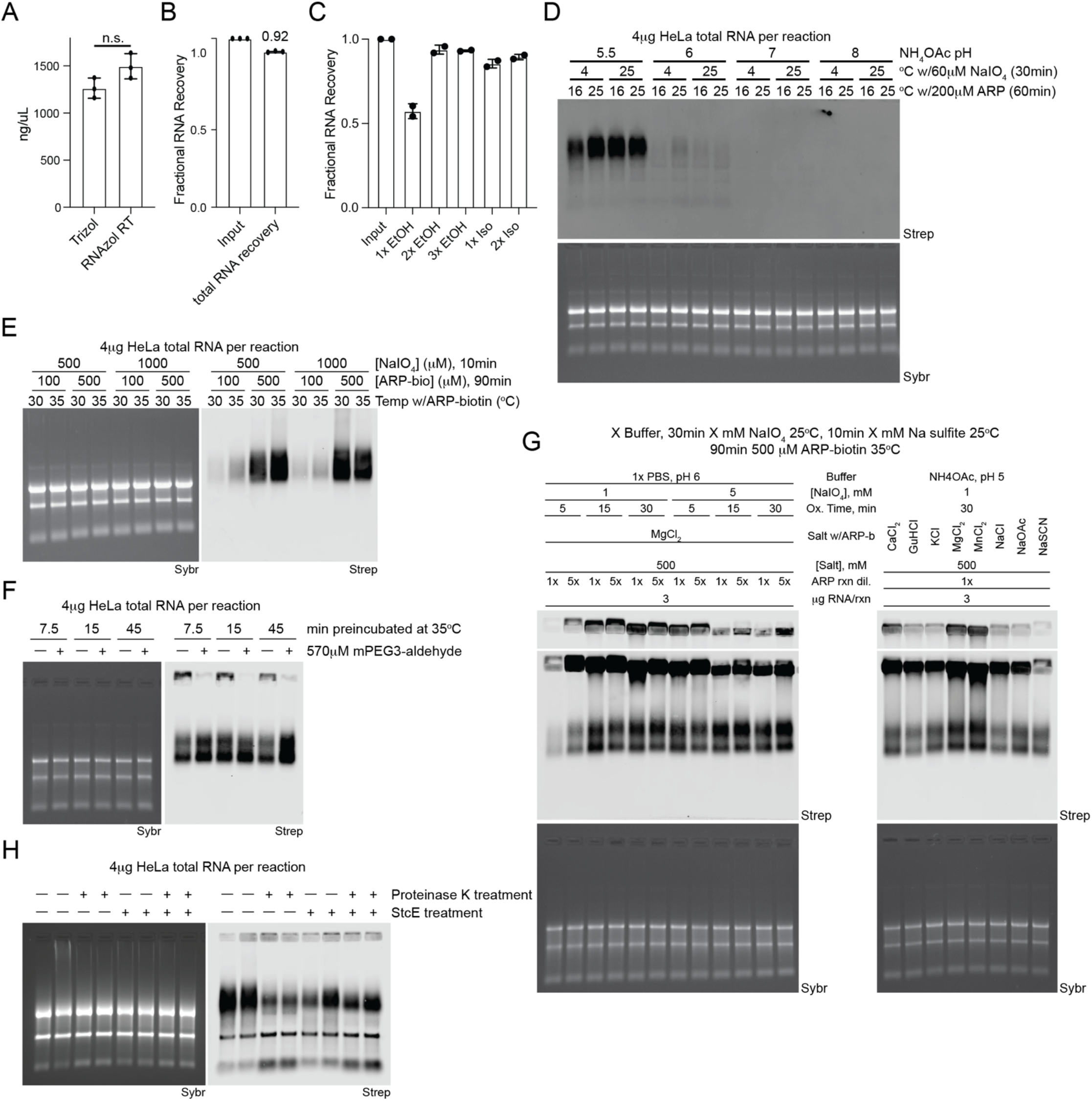
Development of rPAL labeling strategy. **(A)** Nanodrop analysis of total RNA extracted from HeLa cells. Cells were lysed with either Trizol or RNAzol RT and processed as per the manufacturer’s recommendations with RNA cleanups performed using Zymo columns (**Methods**). Each datapoint (biological triplicate) is displayed with the SEM. (**B**) Nanodrop analysis of total RNA extracted from HeLa cells isolated with RNAzol RT. After isolation, samples were quantified and then subjected to a Zymo column clean up after which each sample was quantified again. Each datapoint (biological triplicate) is displayed with the SEM. (**C**) Nanodrop analysis of small RNA extracted from HeLa cells isolated with RNAzol RT. After isolation, samples were quantified and then subjected to a Zymo column clean up. During the precipitation step, before binding to the Zymo columns, indicated ratios of ethanol or isopropanol were added. After Zymo column cleanup, each sample was quantified again. Each datapoint (biological duplicate) is displayed with the SEM. (**D**) RNA blotting of total RNA from HeLa cells. Total RNA was subjected to various reaction conditions including NH_4_OAc buffer pH changes, the temperature at which diol oxidation occurred, and the temperature at which the aldehyde ligation occurred. In gel detection of total RNA with SybrGold (Sybr, bottom) and on membrane detection of biotin (Streptavidin-IR800, top) is shown. (**E**) Blotting as in D with variations in the concentration of NaIO_4_ used for oxidation and ARP-biotin used for aldehyde ligation as well as the temperature at which the aldehyde ligation occurred at. Sybr and Strep detection is as in D. (**F**) Blotting as in D with evaluation of performing a blocking step before the NaIO_4_ oxidation with mPEG3-aldehyde for the indicated times at 35°C. Sybr and Strep detection is as in D. (**G**) Blotting as in D with evaluation of buffer conditions, NaIO_4_ concentration, ARP-biotin concentrations, and oxidation times for generating specific rPAL signal. Sybr and Strep detection is as in D. (**H**) Blotting as in D with evaluation of enzymatic digestions of the RNA samples after extraction using RNAzol RT. RNA was either not digested or subjected to digestion with Proteinase K (45 minutes at 37°C), StcE (35 minutes at 37°C), or both enzymes (sequentially). Sybr and Strep detection is as in D.

**Fig. S2.**
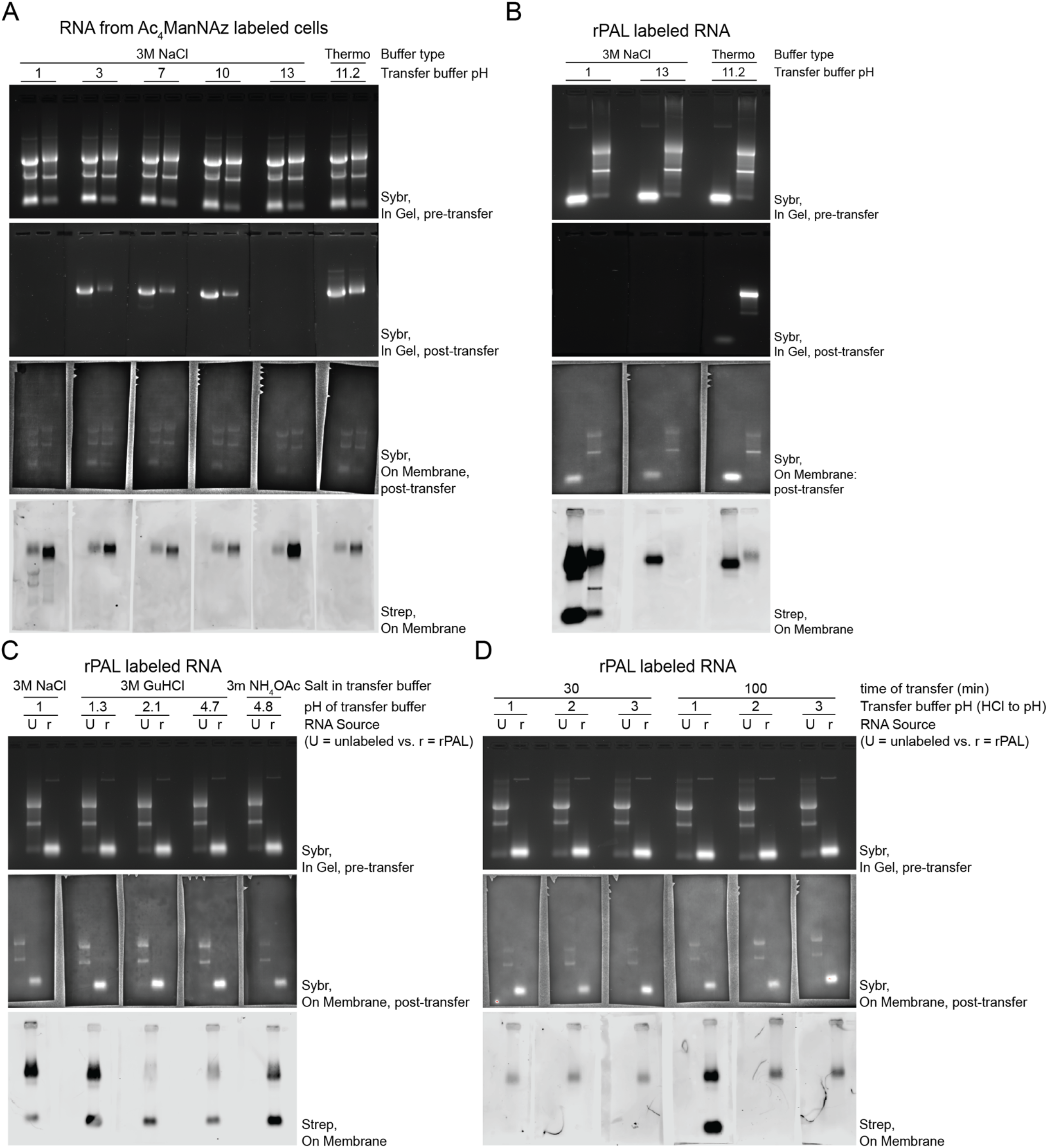
Optimization of RNA Northern blotting transfer conditions. (**A**) RNA northern blot transfer optimization. Small and total RNA samples from HeLa cells labeled with Ac_4_ManNAz and visualized with copper-free click of DBCO-biotin were electrophoresed in a denaturing agarose gel and imaged with SybrGold (top row), after transferring with buffers of various pHs the gel was again imaged (second row) as well as the membrane where the RNA was transferred to (third row). Finally, sialoglycoRNAs were visualized with streptavidin-IR800 (Strep, bottom row). (**B**) RNA northern blot transfer optimization as in A, assessing high and low pH conditions for sialoglycoRNAs transfer detecting with rPAL labeling. (**C**) RNA northern blot transfer optimization as in B, screening buffer salt composition as well as pH. (**D**) RNA northern blot transfer optimization as in B, assessing the time and pH dependency of transfer efficiency.

**Fig. S3.**
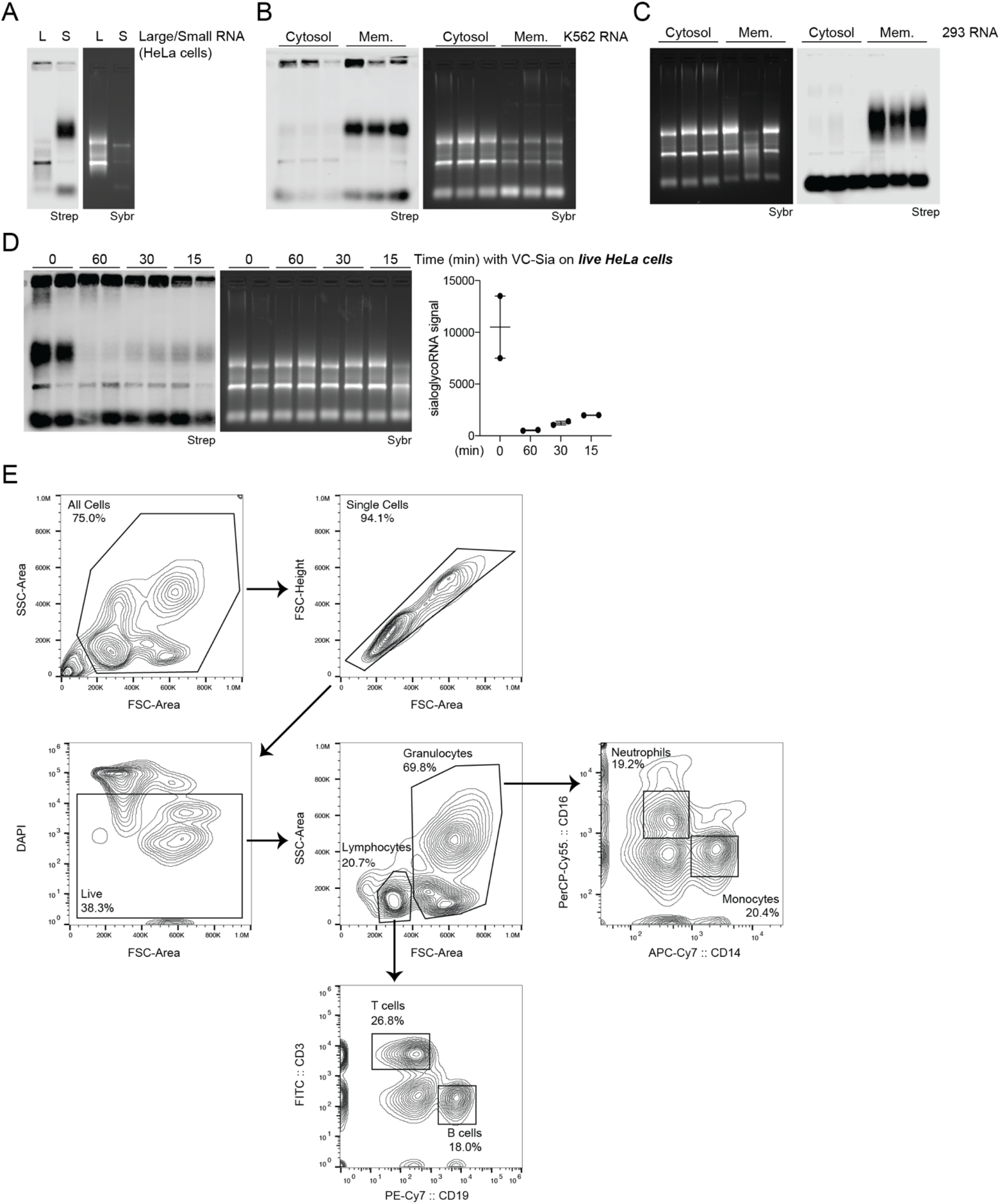
Subcellular localization of rPAL signal and flow cytometry analysis for sorting of human PBMCs. (**A**) RNA blotting of HeLa RNA that after isolation was subjected to a size fractionation, separating out large (L, > ∼200 nts) from small (S, < ∼200 nts) RNAs. Detection of the resulting sialoglycoRNA signal was accomplished by rPAL labeling, in gel detection of total RNA with SybrGold (Sybr, bottom) and on membrane detection of biotin (Streptavidin-IR800, top) is shown. (**B**) RNA blotting of RNA that was isolated from HEK293 cells fractionated for their crude membranes. The soluble cytosol is a fraction from this procedure that serves as a non-membrane enriched control. (**C**) RNA blotting of RNA that was isolated from K562 cells fractionated for their crude membranes. Sybr and Strep detection is as in B. (**D**) RNA blotting of total RNA from HeLa cells that prior to RNAzol RT lysis were exposed to VC-Sialidase treatment in the culture media for the indicated times. After treatment, cells were washed twice with 1x PBS, and then RNA collected and labeled with rPAL. Sybr and Strep detection is as in A. SialoglycoRNA signal was quantified and plotted (right). Each datapoint (biological duplicate) is displayed with the SEM. (**E**) Scatter plots of one of three human peripheral blood mononuclear cell (PBMC) sorting experiments. Each plot represents serial gates drawing to isolate live single cells and further fractionation CD19 (B cells), CD3 (T cells), CD14 (monocytes and macrophages), and CD16 (NK cells, monocytes, and neutrophils). These four populations were then sorted into FACS Buffer for later RNA extraction and sialoglycoRNA analysis. Arrows denote the gating scheme used to isolate the four final populations of cells.

**Figure S4.**
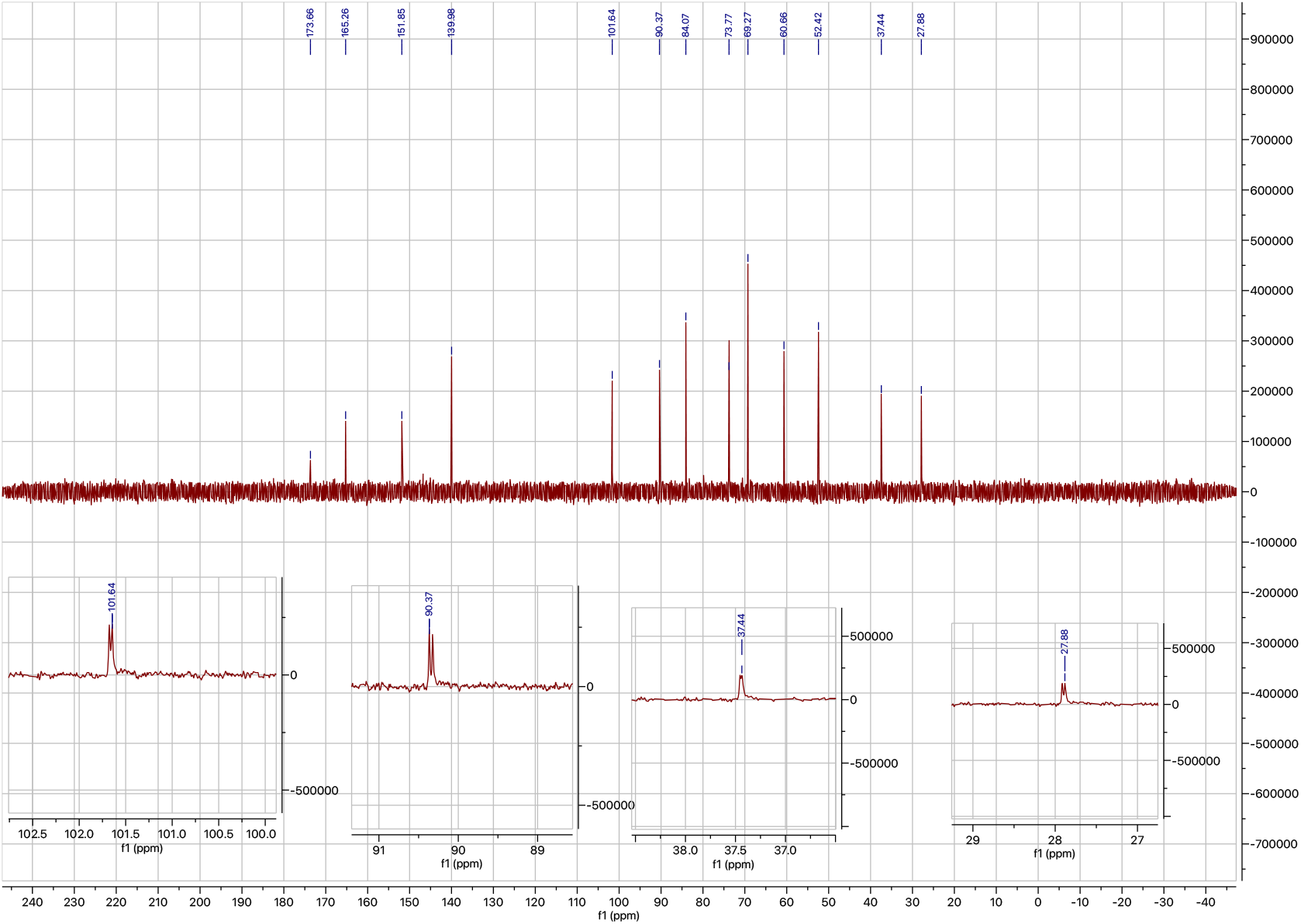
^13^C NMR spectrum of commercial **acp3U** with expanded regions.

**Figure S5.**
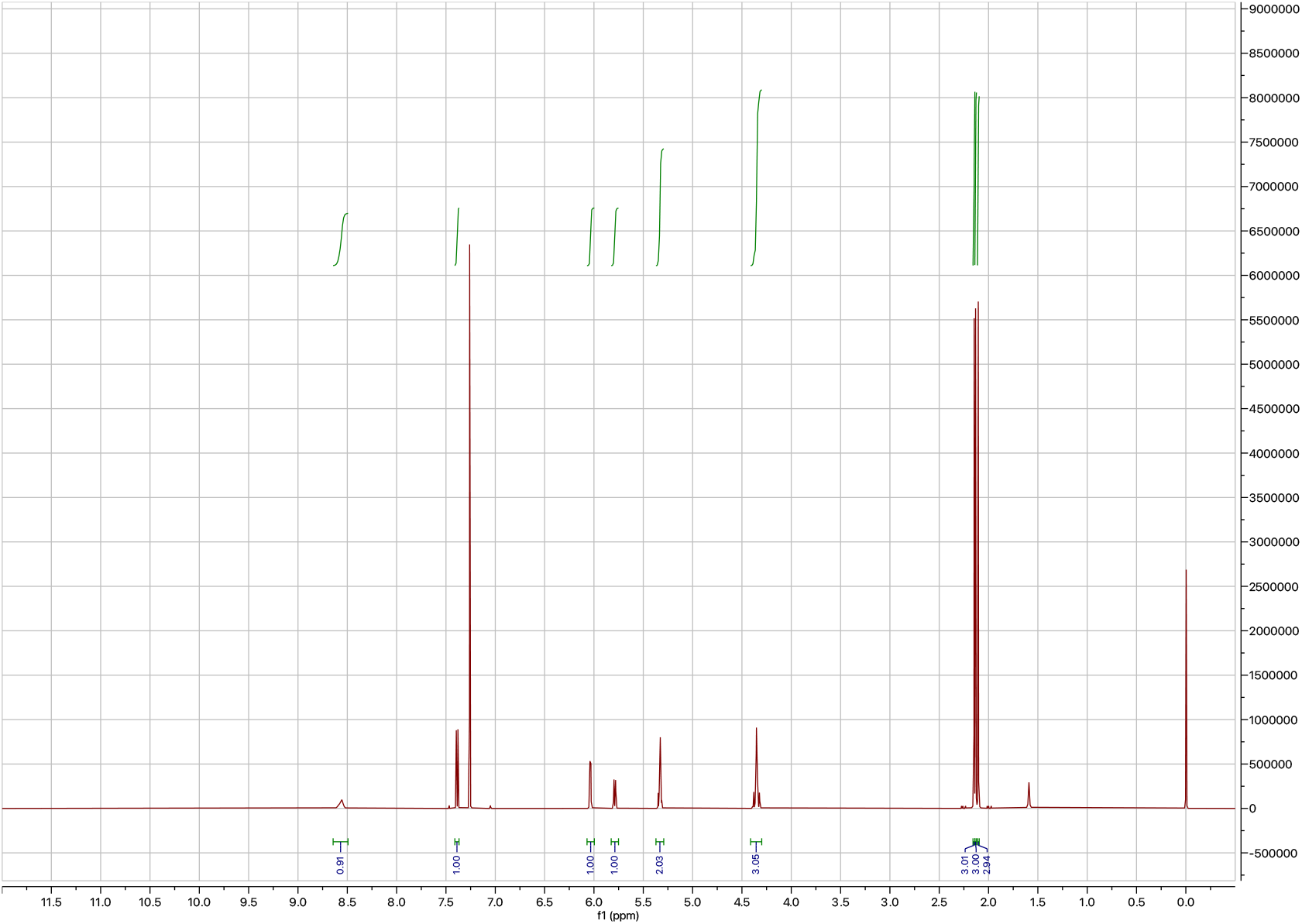
^1^H NMR spectrum of **S1**.

**Figure S6.**
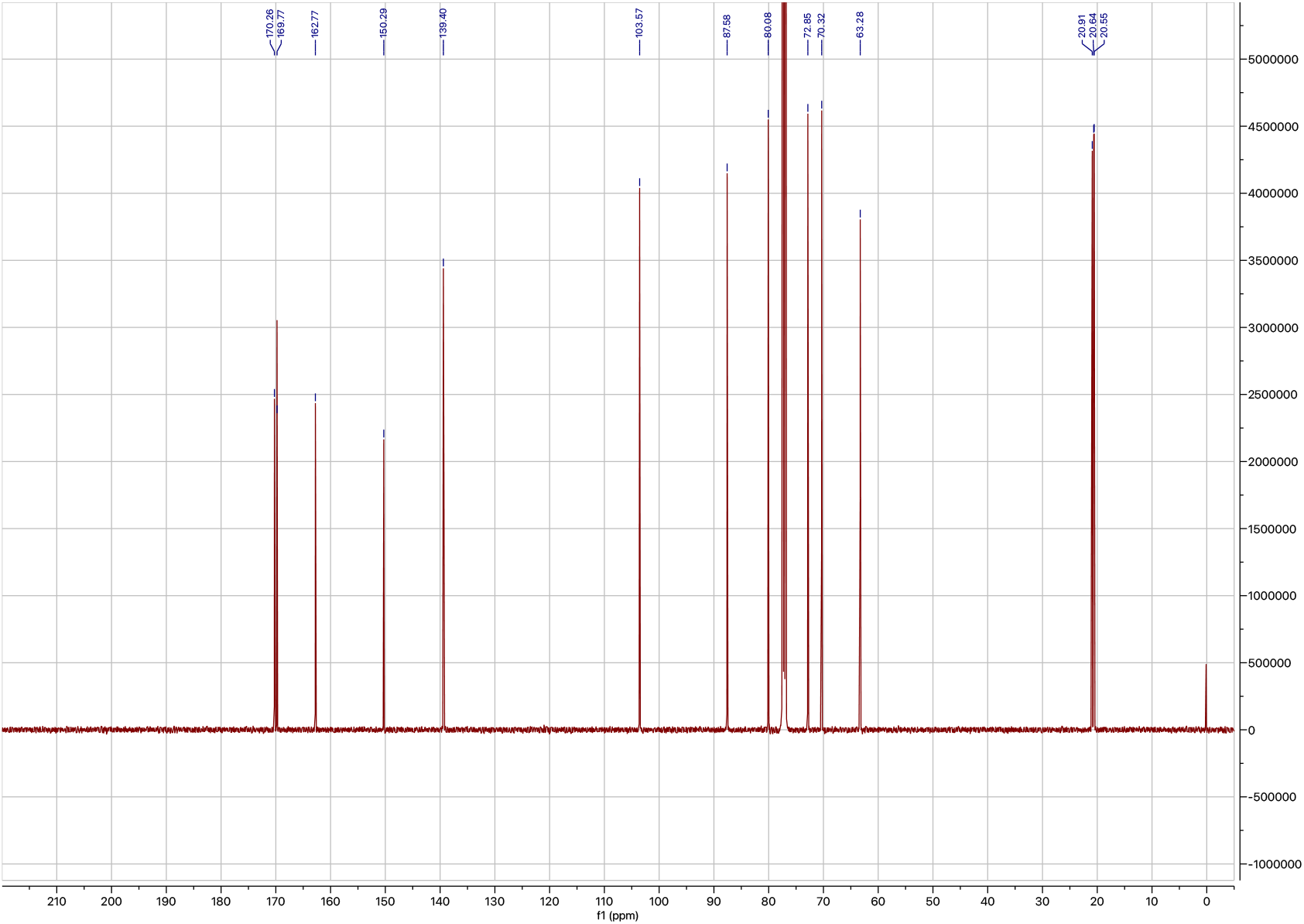
^13^C NMR spectrum of **S1**.

**Figure S7.**
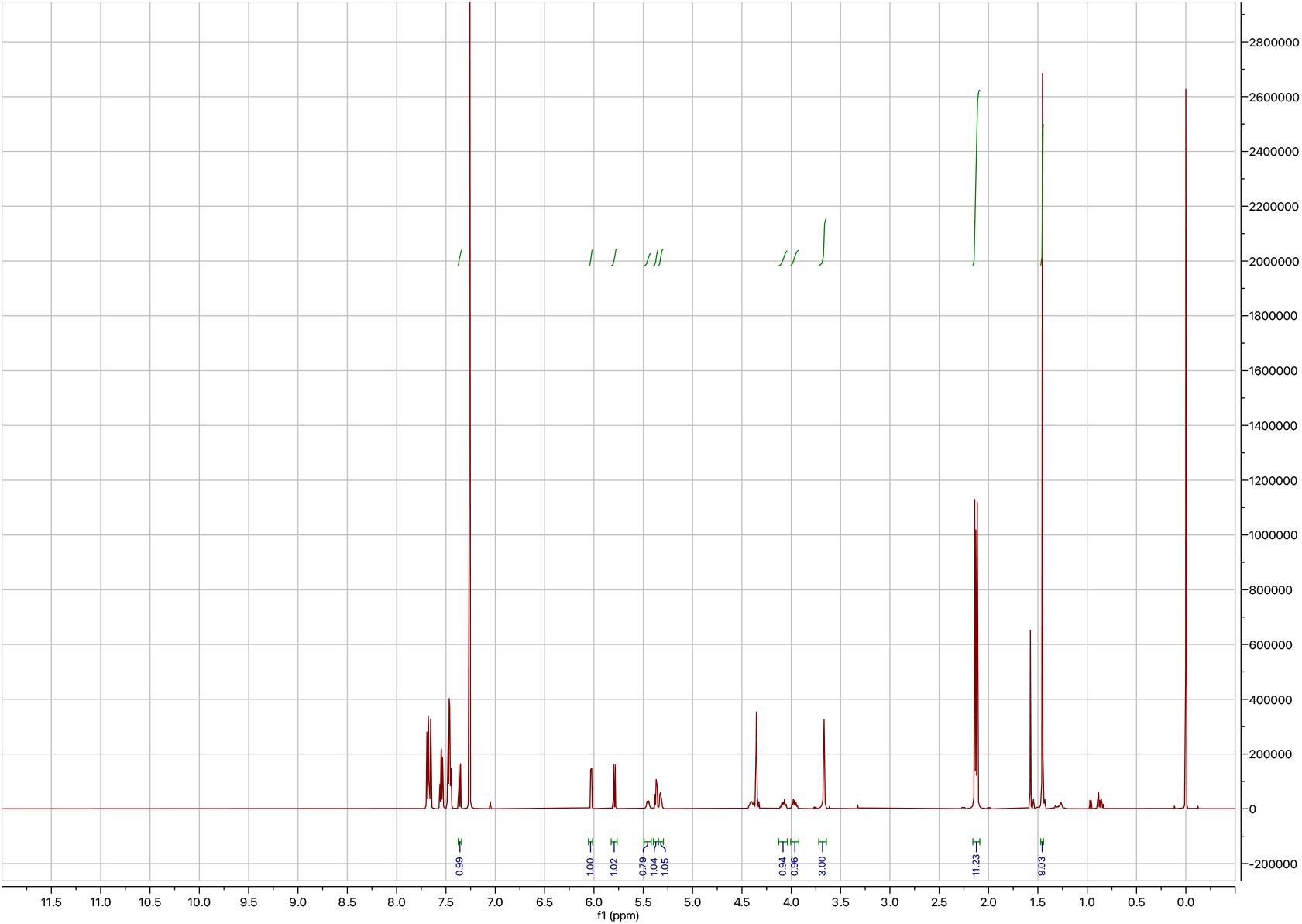
H NMR spectrum of **S2** + PPh_3_O.

**Figure S8.**
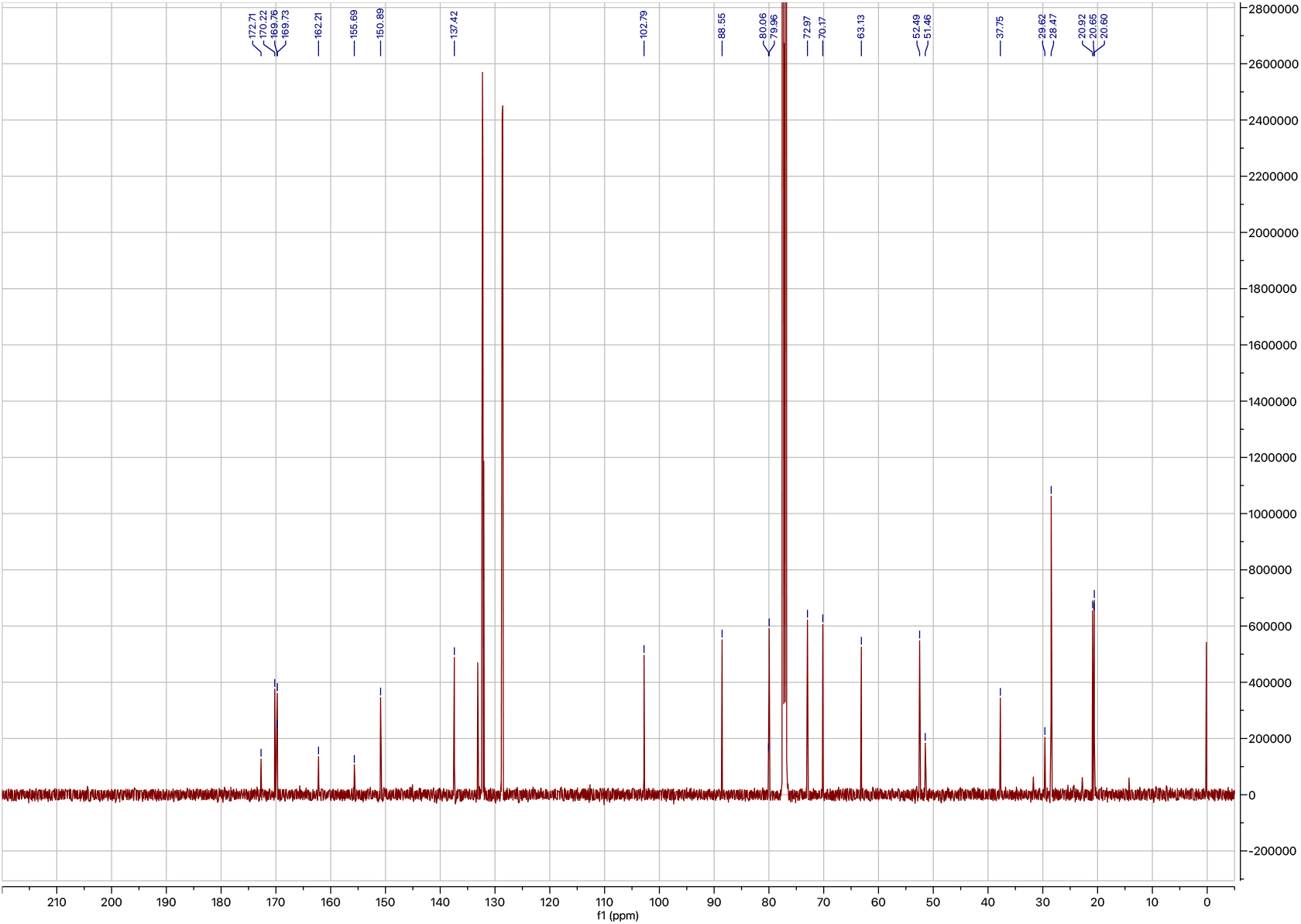
^13^C NMR spectrum of **S2** + PPh_3_O.

**Figure S9.**
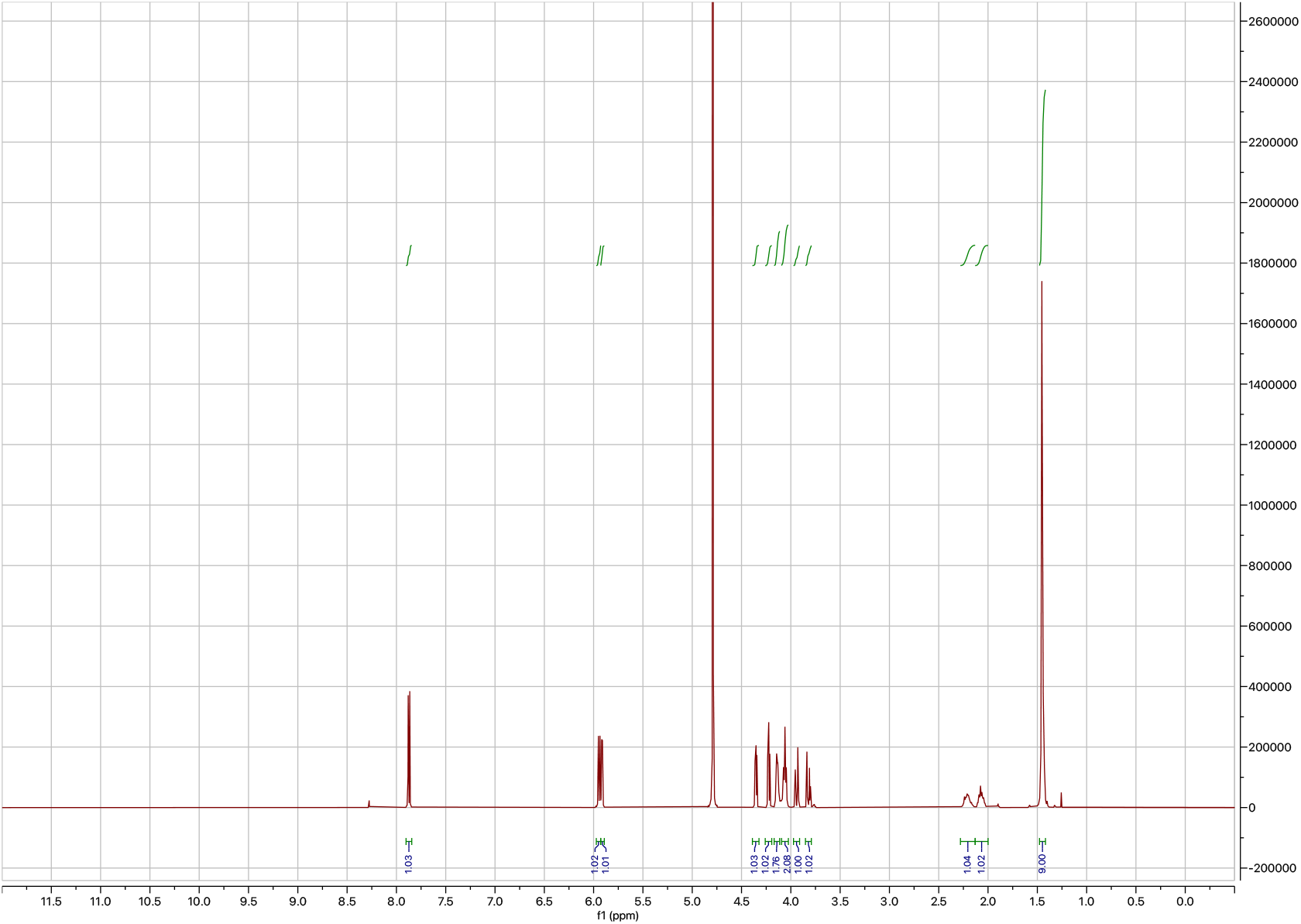
^1^H NMR spectrum of **S3**.

**Figure S10.**
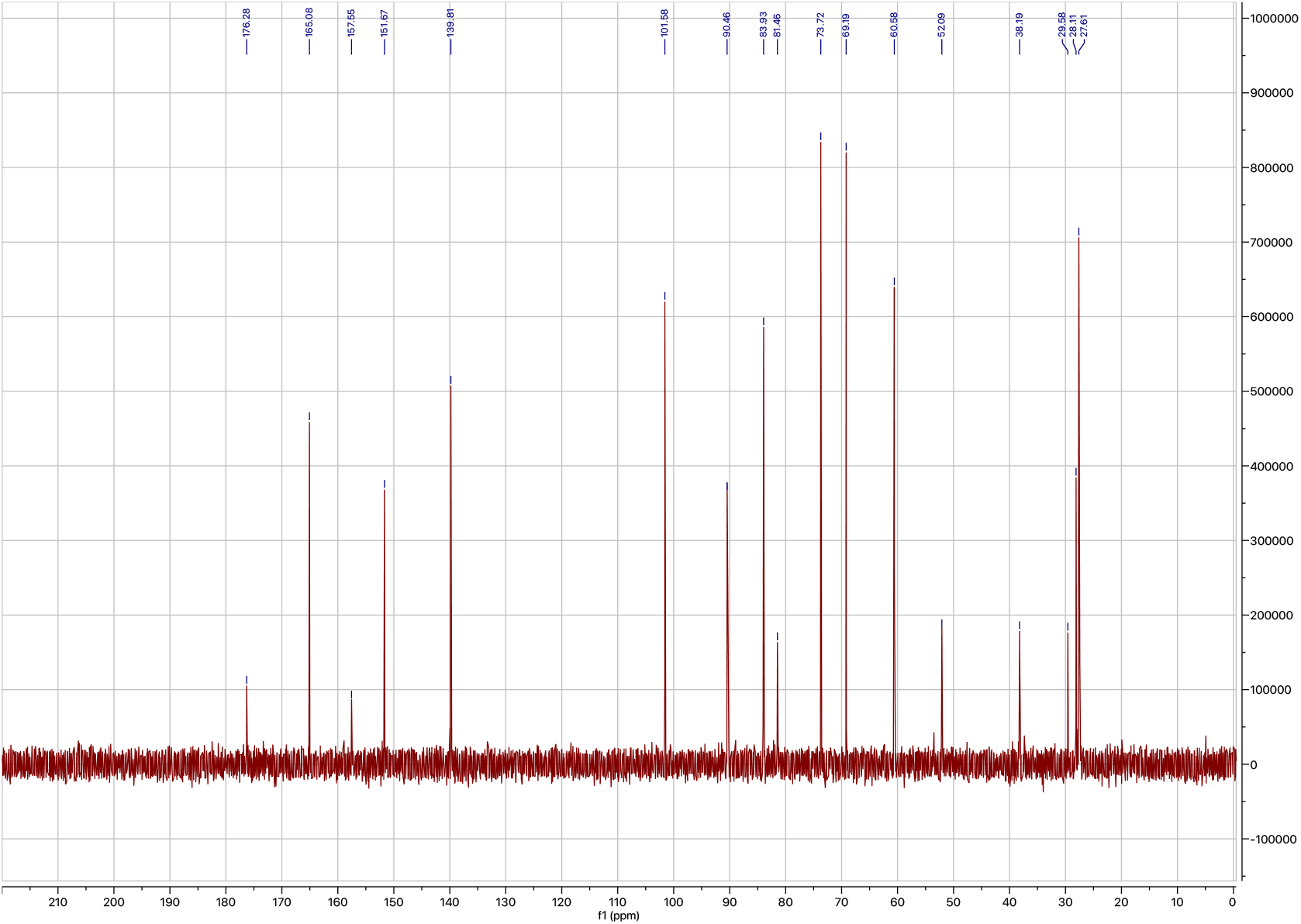
^13^C NMR spectrum of **S3**.

**Figure S11.**
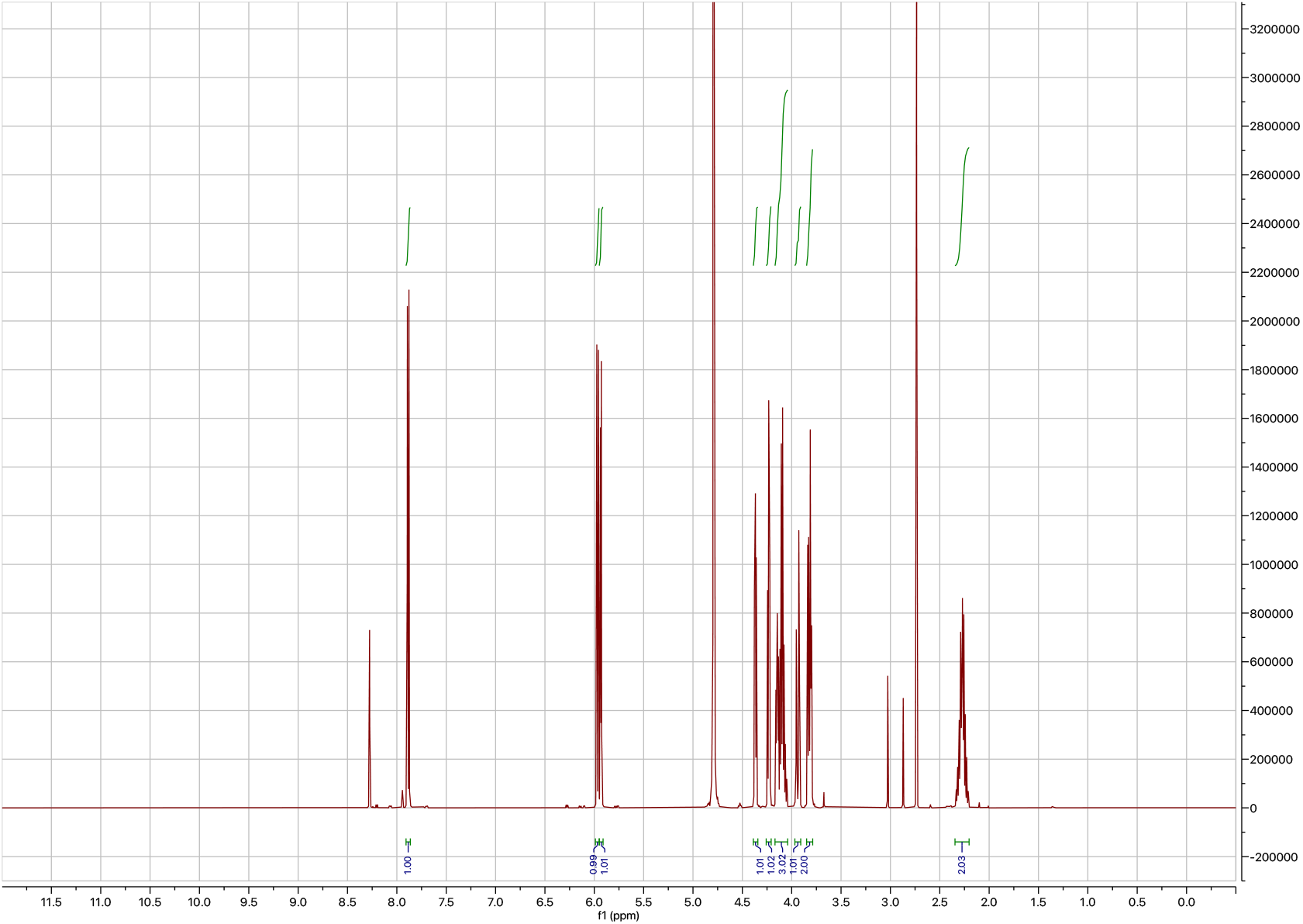
^1^H NMR spectrum of **acp3U**.

**Figure S12.**
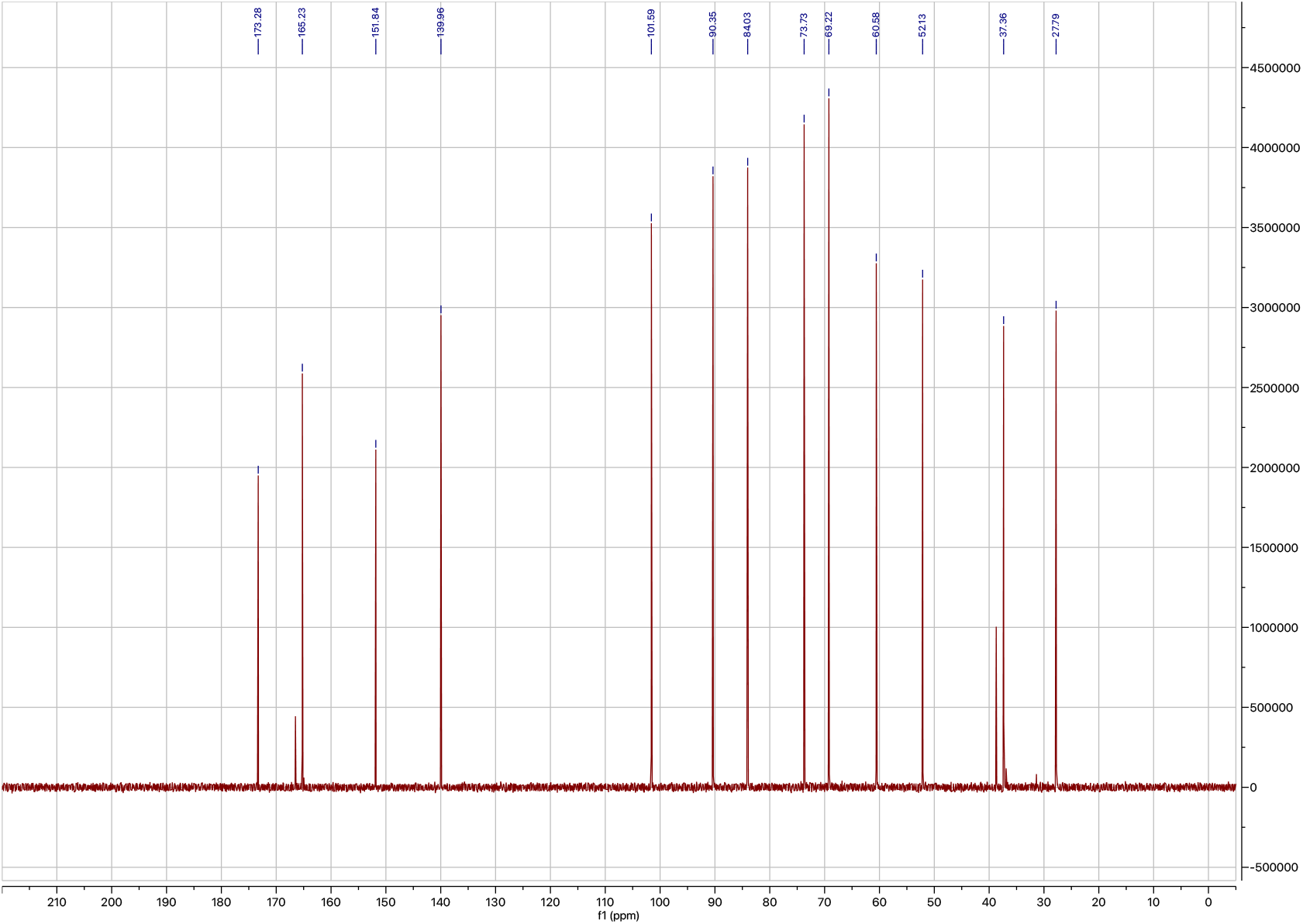
^13^C NMR spectrum of **acp3U**.

**Figure S13.**
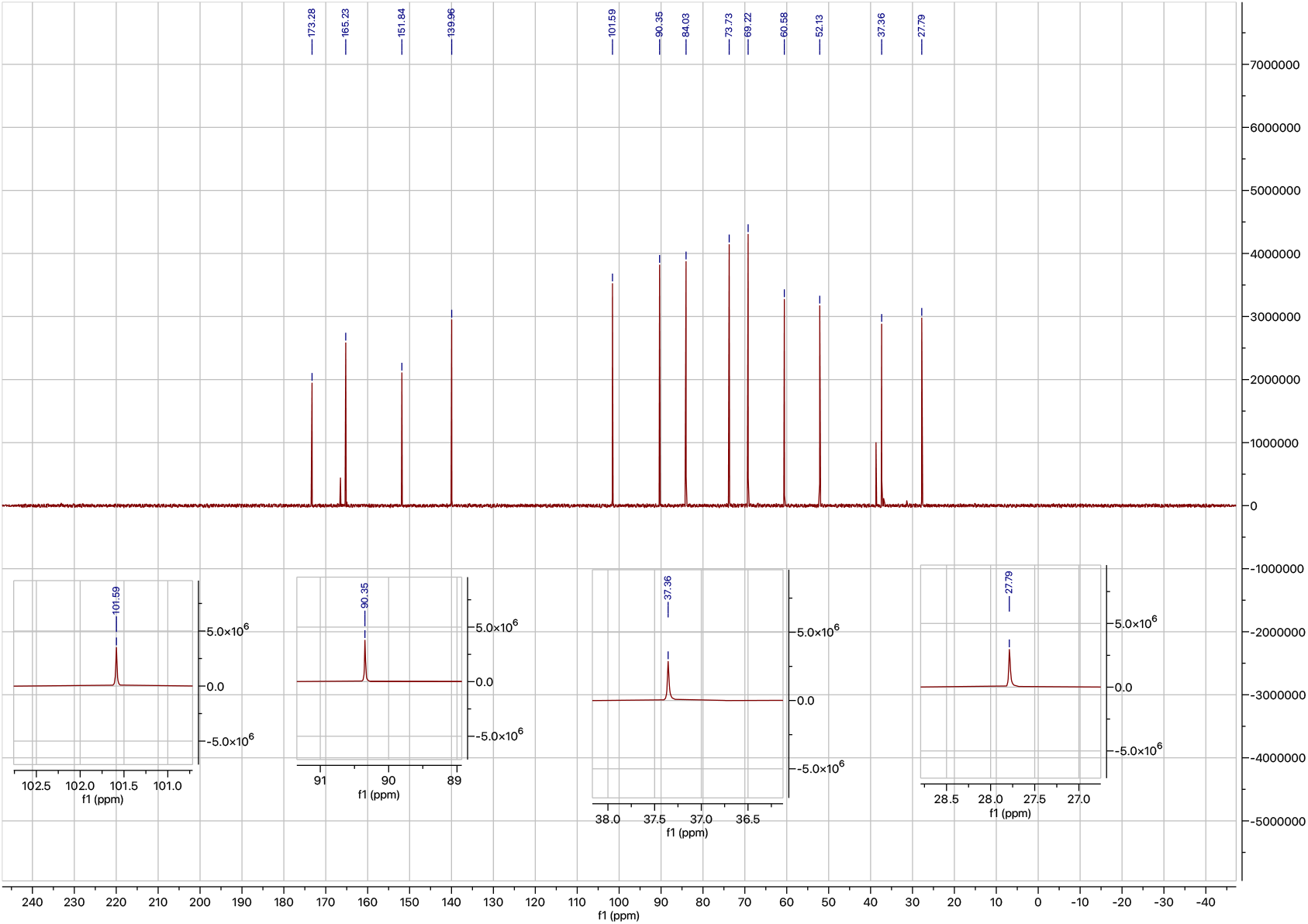
^13^C NMR spectrum of synthetic **acp3U** with expanded regions.

**Figure S14.**
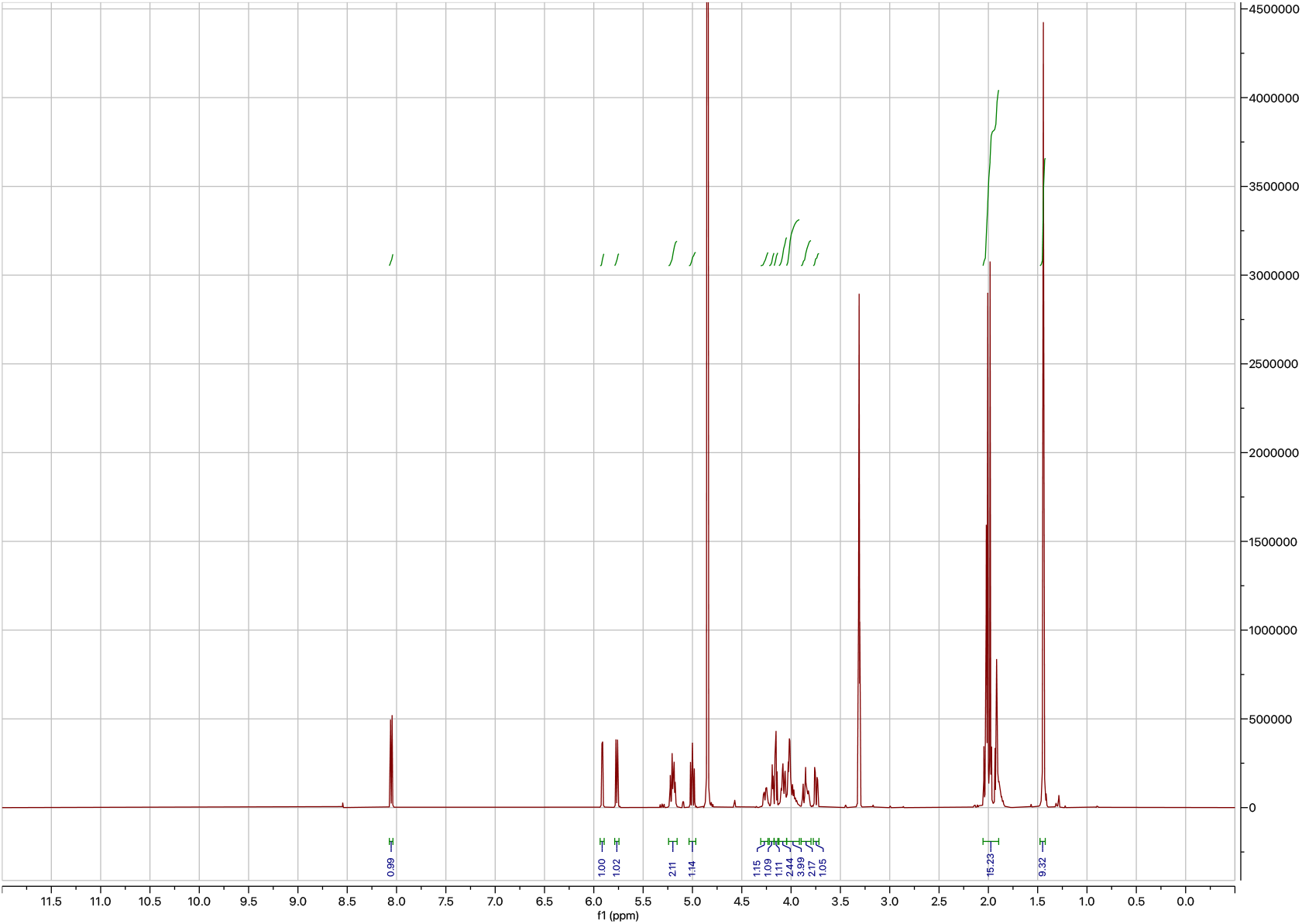
^1^H NMR spectrum of **S4**.

**Figure S15.**
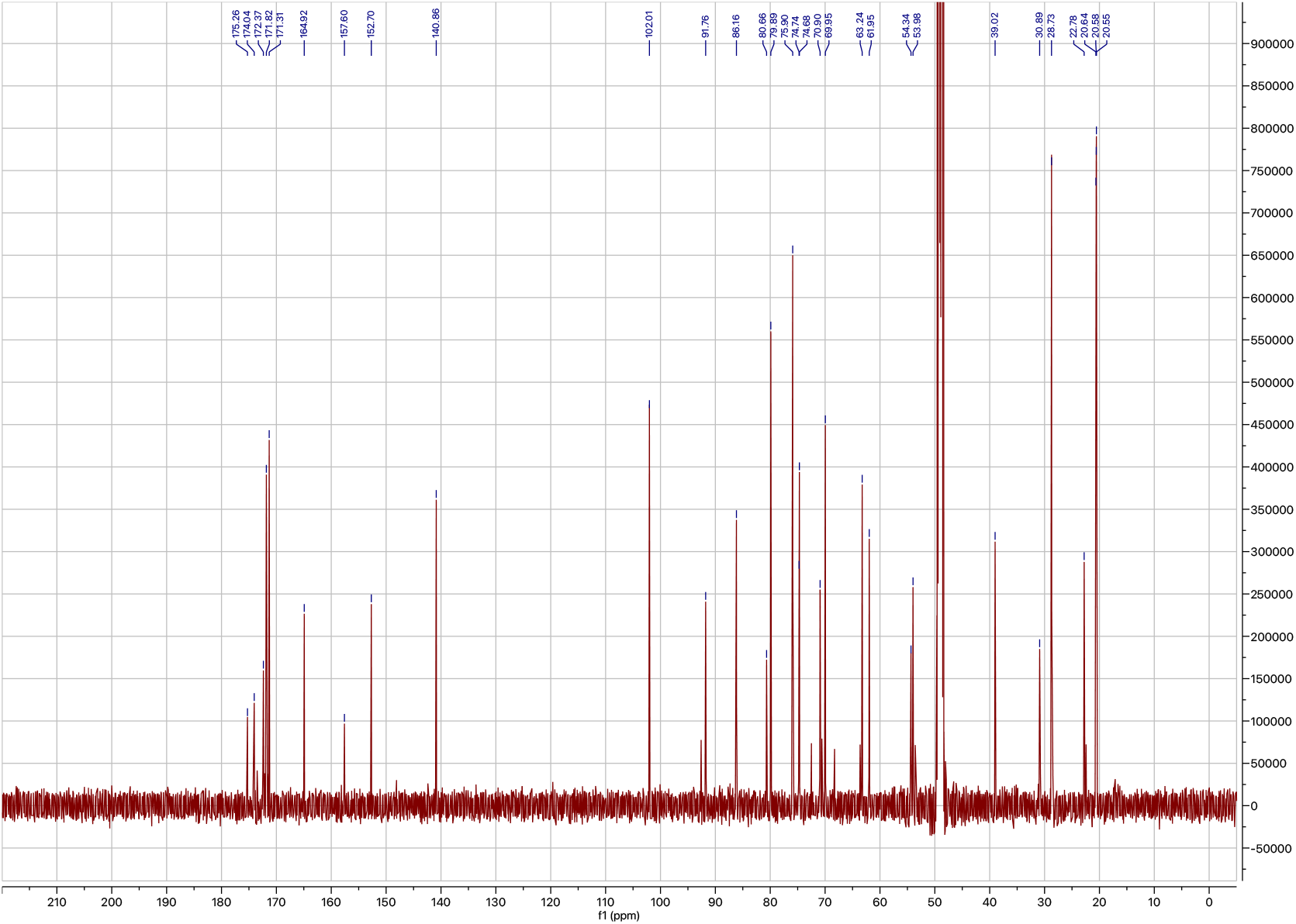
^13^C NMR spectrum of **S4**.

**Figure S16.**
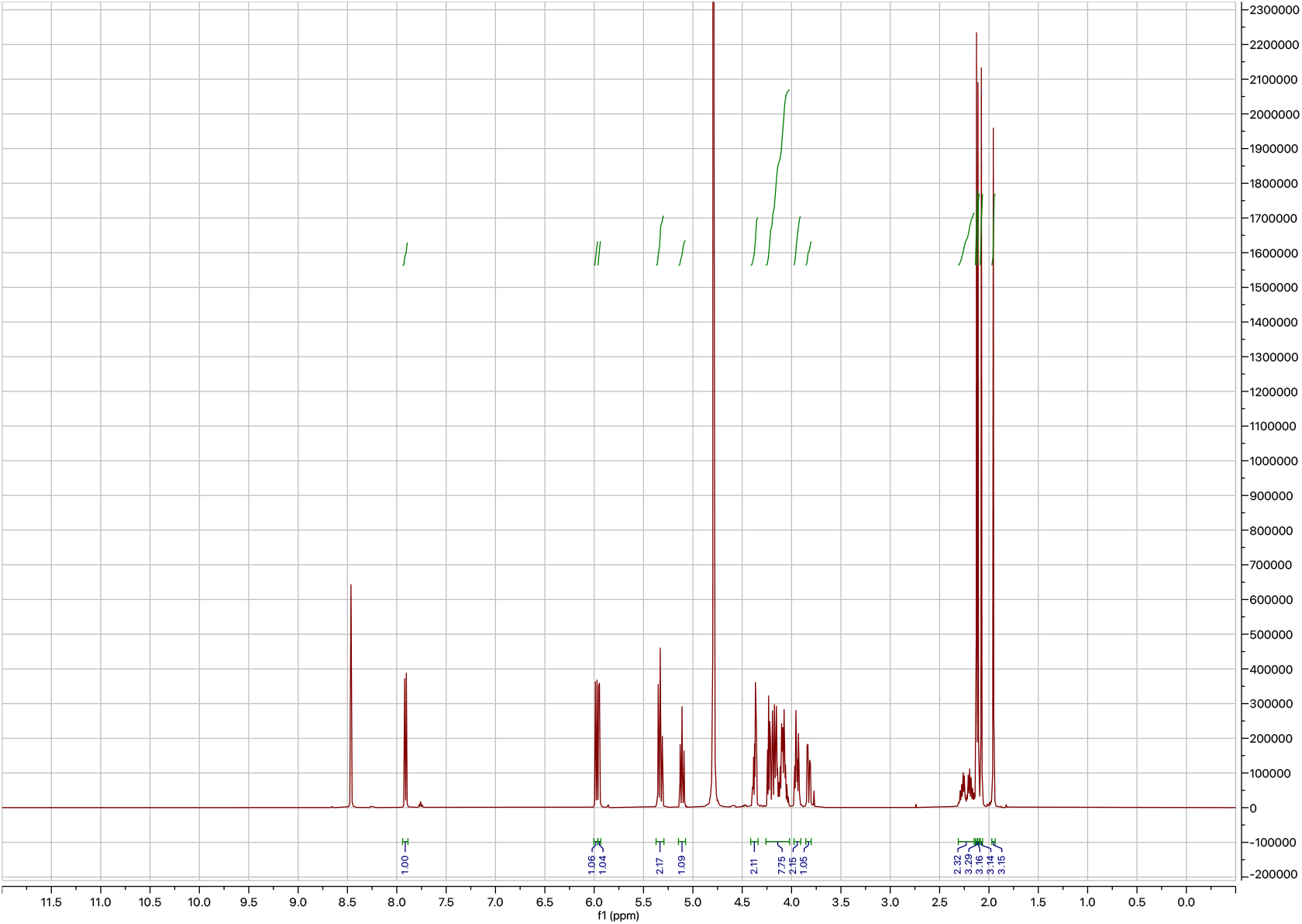
^1^H NMR spectrum of **S5**.

**Figure S17.**
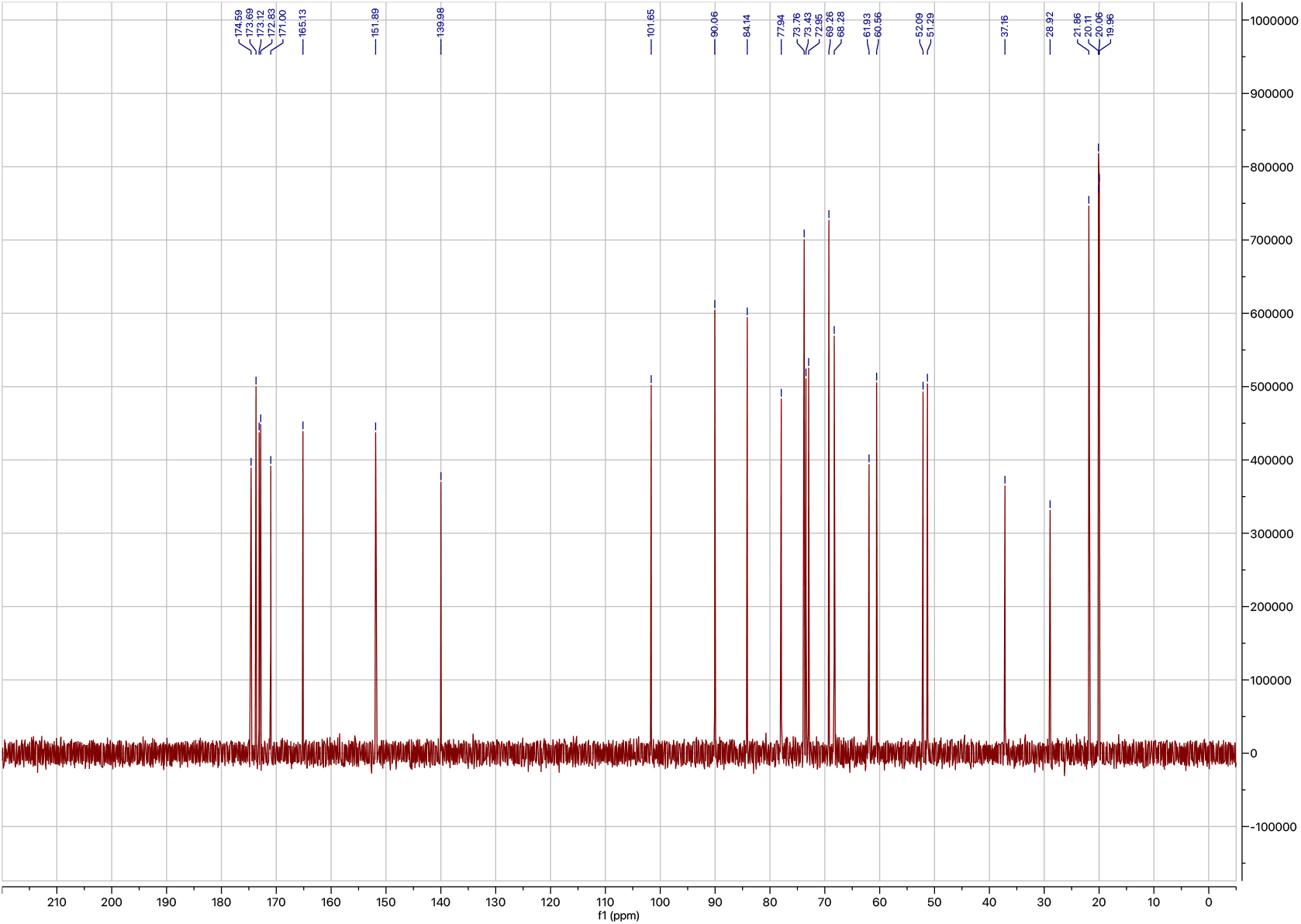
^13^C NMR spectrum of **S5**.

**Figure S18.**
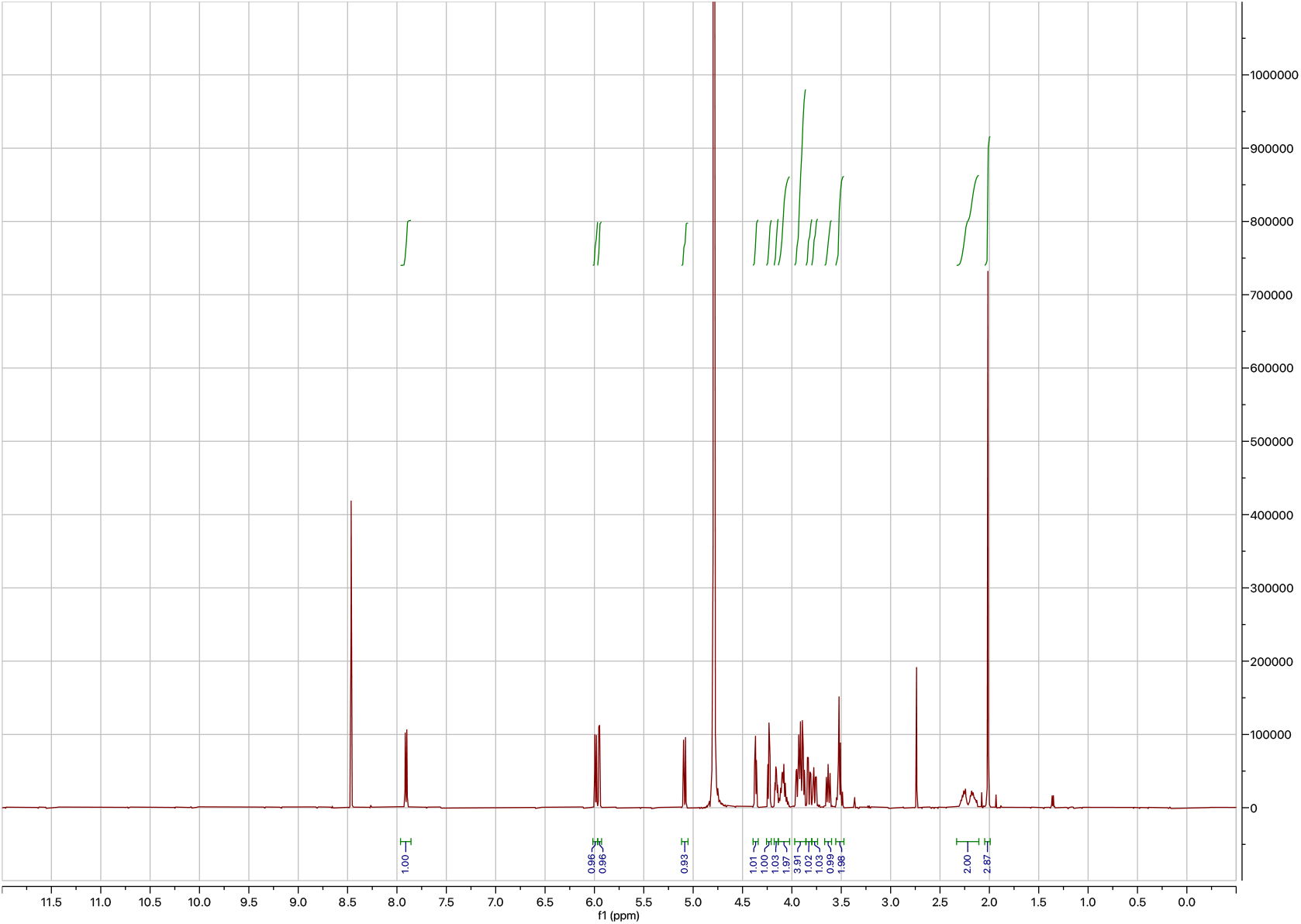
^1^H NMR spectrum of **acp3U-GlcNAc**.

**Figure S19.**
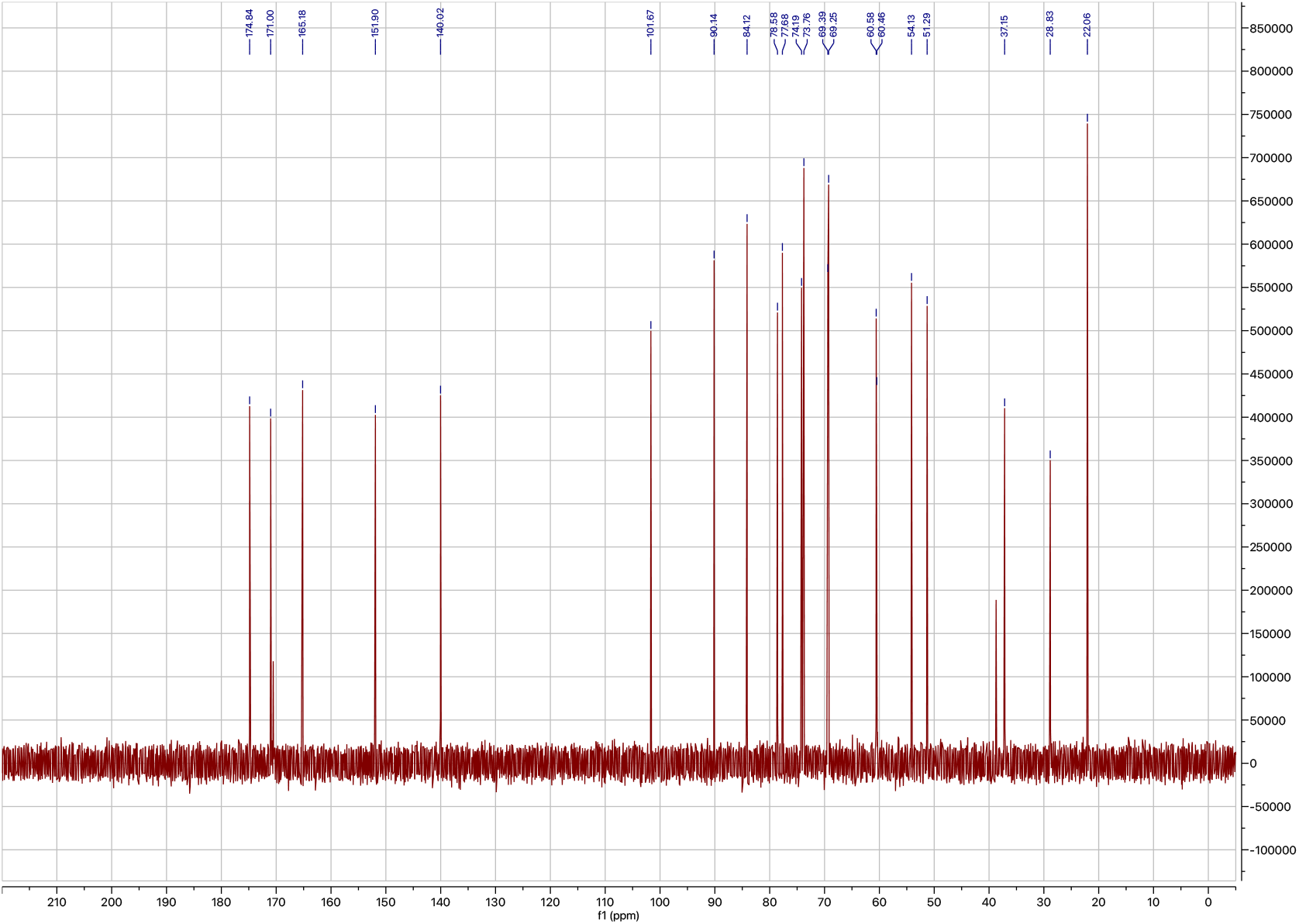
^13^C NMR spectrum of **acp3U-GlcNAc**.

**Fig. S20.**
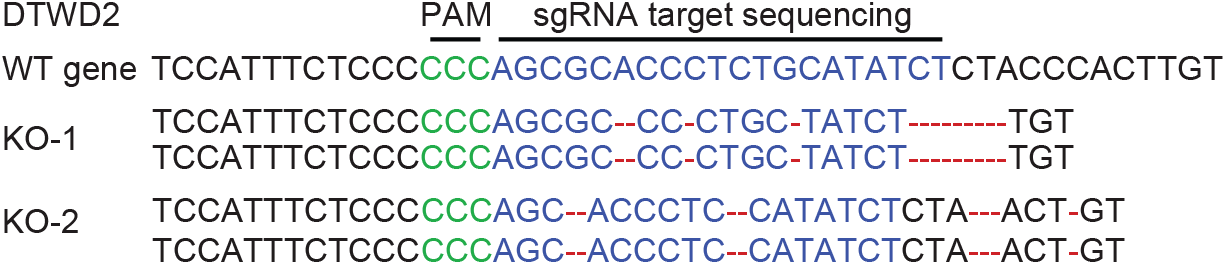
Knockout of DTWD2 in U2OS cells. Sequence alignment of DTWD2 coding sequences targeted for Cas9-mediated cleavage from WT and DTWD2 KO U2OS cell lines. Protospacer adjacent motif (PAM), green; sgRNA target, blue; mutated sequence, red.

